# Inhibition of Indirect Pathway Activity Causes Abnormal Decision-Making In a Mouse Model of Impulse Control Disorder in Parkinson’s Disease

**DOI:** 10.1101/2024.02.19.581062

**Authors:** Xiaowen Zhuang, Julia Lemak, Sadhana Sridhar, Alexandra B. Nelson

## Abstract

Healthy action selection relies on the coordinated activity of striatal direct and indirect pathway neurons. In Parkinson’s disease (PD), in which loss of midbrain dopamine neurons is associated with progressive motor and cognitive deficits, this coordination is disrupted. Dopamine replacement therapy can remediate motor symptoms, but can also cause impulse control disorder (ICD), which is characterized by pathological gambling, hypersexuality, and/or compulsive shopping. The cellular and circuit mechanisms of ICD remain unknown. Here we developed a mouse model of PD/ICD, in which ICD-like behavior was assayed with a delay discounting task. We found that in parkinsonian mice, the dopamine agonist pramipexole drove more pronounced delay discounting, as well as disrupted firing in both direct and indirect pathway neurons. We found that chemogenetic inhibition of indirect pathway neurons in parkinsonian mice drove similar phenotypes. Together, these findings provide a new mouse model and insights into ICD pathophysiology.

## INTRODUCTION

We often weigh immediate versus distant costs and benefits in making decisions. Impulsive decision-making is characterized by intolerance for long-term costs and preference for more immediate rewards. Impulsivity is seen in a number of neuropsychiatric conditions, including neurodegenerative disorders, psychiatric disease, and drug addiction ^1,2^. One notable example is impulse control disorder (ICD), a complication of Parkinson’s disease (PD) treatment. PD is characterized by progressive degeneration of midbrain dopamine neurons, which contributes to motor impairment, such as slowing of movement (bradykinesia), tremor and rigidity ^3^. Dopamine replacement therapy, particularly with D2/3-type receptor (D2/3R) agonists, alleviates motor deficits, but can be complicated by the development of ICD. In response to dopamine agonists, up to 40% of PD patients develop non-motor symptoms like pathological gambling, binge eating, or hypersexuality; this cognitive-behavioral syndrome is termed ICD ^4,5^. Our current understanding of ICD is primarily informed by epidemiological and imaging studies in clinical populations; there are few studies in animal models, and the pathophysiological mechanisms remain unknown ^6-9^.

The cognitive profile of PD/ICD provides a few clues as to its origins. Those with ICD show a preference for immediate rewards, and an intolerance for delays. This has been studied by measuring delay discounting, a normal cognitive phenomenon in which the value of a reward decreases according to the time needed to wait for it. Notably, those with ICD are more likely to choose an immediate but small reward over a delayed/large reward in delay discounting tasks ^10-12^. Previous work suggests that delay discounting is mediated in large part by the frontal cortex and the striatum (caudate and putamen), as well as by dopamine ^13,14^. In healthy nonhuman primates, dopamine agonist infusion in the striatum induced impulsive choices in monkeys during a delay discounting task ^15^. More specifically, activity in striatal neurons encodes key variables of delay discounting behavior ^16-18^.

One of the most distinctive features of ICD in PD is its relationship to dopamine agonist medication. Reducing the dose or discontinuing the dopamine agonist typically eliminates symptoms of ICD ^19^. This implies that dopamine signaling may lead to a reversible change in neural activity, potentially within corticostriatal circuits, which drives ICD. Within the striatum, dopamine regulates striatal projection neurons, medium spiny neurons (MSNs). Direct pathway MSNs (dMSNs) express the dopamine D1 receptor (D1R) and indirect pathway MSNs (iMSNs) express the D2 receptor (D2R) ^20^. Striatal dopamine release is hypothesized to excite dMSNs and inhibit iMSNs, based on *ex vivo* and *in vivo* recordings ^21-25^. Indeed, in mouse models of PD, treatment with the dopamine precursor levodopa, or dopamine agonists, causes acute bidirectional changes in dMSN and iMSN activity ^24,26^. Another complication of dopamine replacement therapy, levodopa-induced dyskinesia (LID) is associated with especially high dMSN activity and low iMSN activity ^24,27,28^. While the neural correlates of ICD are unknown, one possibility is that chronic dopamine depletion in PD leads to circuit vulnerability, and in this context dopamine agonists trigger an imbalance in dMSN and iMSN activity driving ICD.

To address these questions, we created a mouse model of PD/ICD. In mildly parkinsonian (but not in healthy) mice, the ICD-associated dopamine agonist pramipexole (PPX) led to alterations in delay discounting behavior reminiscent of those seen in PD/ICD ^4,5^. We used this model to explore the cellular and circuit mechanisms of ICD. We found that chemogenetic inhibition of iMSNs in the cognitive region of the striatum drove impulsive decision-making. We also found that PPX induced marked bidirectional changes in dMSN and iMSN firing in parkinsonian mice. Chronic PPX treatment further potentiated these changes in striatal physiology and decision-making behavior. Taken together, our findings provide a robust mouse model of ICD, and shed light on how dopaminergic agonists may induce pathological impulsivity in PD.

## RESULTS

### Treatment with the dopamine agonist pramipexole causes impulsive decision-making in parkinsonian mice

To model early-stage Parkinson’s disease (PD), for which dopamine agonist therapy is often used ^35^, we injected the dopaminergic neurotoxin 6-OHDA bilaterally in the dorsolateral striatum (DLS). This approach resulted in partial loss of midbrain dopamine neurons, with greater impact on axons in the dorsal striatum (Fig. S1A). Using tyrosine hydroxylase (TH) as a surrogate marker for dopamine neurons, we found approximately 50% loss of TH signal in the dorsal striatum, with less marked depletion in the ventral striatum (VS; Fig. S1B, see statistics in Figure Legend and Table 1). Dopaminergic cell bodies in the substantia nigra pars compacta (SNc) were also markedly reduced in 6-OHDA-treated versus control mice (Fig. S1C,D). 6-OHDA-treated mice showed mild motor impairment on the accelerating rotarod test (Fig. S1E), consistent with a mild-moderate parkinsonian phenotype. As in people with early-stage PD, motor performance was remediated by treatment with the dopamine D2/3-type agonist, pramipexole (PPX, 0.5 mg/kg; Fig. S1E). Consistent with findings in healthy rodents ^32^, PPX caused an acute reduction in movement in both control and parkinsonian mice. However, increased locomotor activity was seen at later time points in parkinsonian mice, consistent with a therapeutic response (Fig. S1F&G). These findings indicate the bilateral/partial 6-OHDA model shows key behavioral features of early-stage PD, which are responsive to dopamine agonist medication.

To model alterations in decision making seen in impulse control disorder (ICD), we took advantage of a normal cognitive phenomenon, delay discounting, in which the value of a reward is discounted by the time needed to wait for it ^33^. Delay discounting behavior is abnormal in individuals with ICD, with more pronounced discounting, or intolerance for delays ^10-12^. We adapted a rodent delay discounting task for use in healthy and parkinsonian mice (Fig. 1A). Prior to training in the delay discounting task, control and parkinsonian mice underwent behavioral shaping, with two phases of instrumental learning (Fig. 1B, S1H,K). In both phases, parkinsonian mice showed slightly slower response latencies and learning rates, but eventually achieved similar performance (Fig. S1I-M). These results indicate that while the bilateral/partial 6-OHDA model shows mild motor deficits, it does not impair the fundamental capacity for instrumental learning.

**Figure 1.**
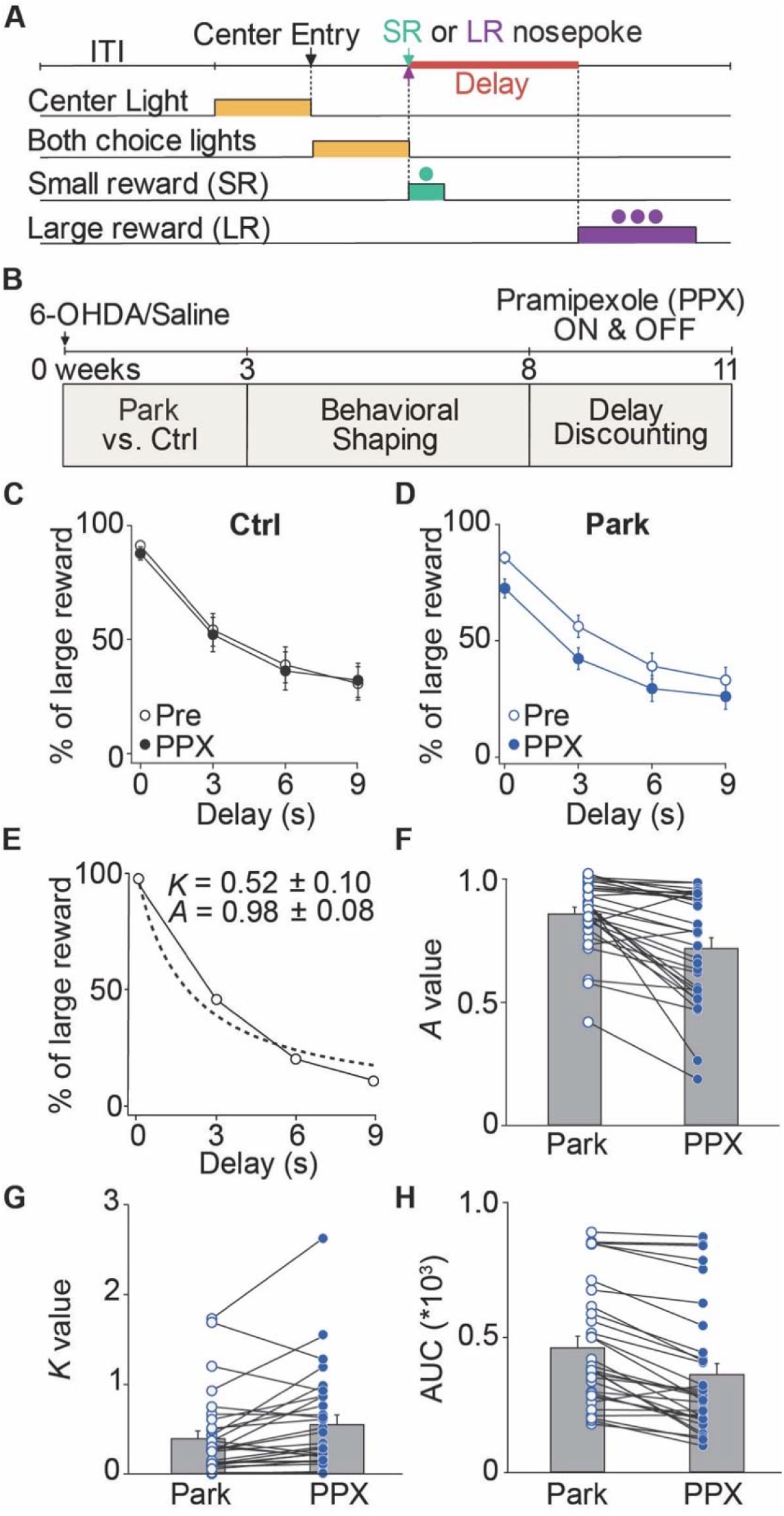
Treatment with the dopamine agonist pramipexole causes impulsive decision-making in parkinsonian mice. (**A**) Delay Discounting task structure. Each delay was tested in a separate block, up to 9 seconds. ITI = inter-trial interval. (**B**) Experimental timeline. (**C, D**) Percentage of trials in which mice chose the delayed/large reward across delays during baseline (open circles) and following pramipexole (PPX) injections (filled circles). PPX did not significantly change decision-making in healthy control mice (C, N = 16; p > 0.05 at all delays), but reduced the likelihood of delayed/large choices at every delay in parkinsonian mice (D, N = 31; 0s: p < 0.001, 3s: p < 0.001. 6s: p < 0.001, 9s: p < 0.01). (**E**) Hyperbolic discounting function, fitted (dashed line) to representative delay discounting behavior in one mouse. The intercept and steepness of the curve were quantified by *A* and *K*, respectively. (**F-H**). *A, K* and area-under-the-curve (AUC) values associated with delay discounting during before (Park) and following PPX treatments (PPX) in parkinsonian mice (N = 31, F, p < 0.001; G, p = 0.01; H, p < 0.001). N, animals. All data presented as means ± SEMs.

We next trained animals in the delay discounting task, during which animals chose between two alternatives: an immediate, small reward, and a larger reward at various delays: 0, 3, 6 and 9 s. During the task, both control and parkinsonian mice showed classic discounting behavior. The likelihood of choosing the large reward declined as the associated delay increased (Fig. 1C,D). There was no significant change in delay discounting behavior between PPX-naïve control and parkinsonian mice. As ICD is associated with dopamine D2/3 agonist treatment in people with PD ^4,5^, we next tested whether PPX altered delay discounting behavior. After baseline sessions, healthy control and parkinsonian mice were tested in PPX sessions (4 h after injection). Consistent with findings in PD patients with ICD ^10-12^, a moderate dose of PPX (0.5 mg/kg) significantly reduced the likelihood of delayed/large reward choices as compared to baseline in the parkinsonian mice (Fig. 1D). Notably, no significant changes were observed in control mice treated with PPX (Fig. 1C), consistent with the lower risk of ICD in people without PD who are treated with PPX ^36^. Together, these findings suggest that like people with PD/ICD, parkinsonian mice are more vulnerable to the effects of PPX on decision-making.

To exclude the possibility that PPX altered discounting behavior indirectly through changes in motivation or attention, we monitored other task outcomes, including omitted trials and response latencies. Mice showed low omission rates during baseline and PPX sessions (Fig. S2A-D). The latency to choose the delayed/large reward progressively increased across delays in both healthy and parkinsonian mice, while the latency to choose the immediate/small reward decreased, until they eventually reached a similar level (Fig. S2E,F). This observation aligns with previous studies suggesting that the anticipation of different reward outcomes modulates the response time in goal-directed behavior ^37,38^. However, in parkinsonian mice treated with PPX, modulation in response latencies by outcome was absent (Fig. S2G), suggesting PPX-induced impairment in goal-directed responding.

To better characterize impulsive decision-making in parkinsonian mice treated with PPX, we fitted a hyperbolic discounting function V = 100\**A*/(1 + *K*D) to each mouse’s delay discounting curve (Fig. 1E). This function has previously been utilized to quantify aspects of discounting behavior ^33,34,39^. The probability of choosing a large reward (V) is devalued by the length of delay (D), scaled by the discounting propensity (*A* and *K*). *K* reflects sensitivity to delays, or an index of the discount rate (steepness of the curve); and parameter *A* reflects sensitivity to reward magnitude (intercept with the y axis) ^40,41^. PPX had variable effects on *A* and *K* in individual parkinsonian mice, but overall led to a decrease in *A* and increase in *K* (Fig 1F&G). The reduction in *A* suggested parkinsonian mice treated with PPX had more difficulty differentiating reward magnitudes in the absence of delay. The increase in *K* (a steeper discounting curve), suggested PPX-treated animals more heavily weighted the cost of delay. We also quantified the shape of the discounting curve by measuring area-under-the-curve (AUC); in this analysis, a decrease in AUC indicated an in increase in impulsive choice ^42,43^. PPX treatment led to a significant reduction in AUC in parkinsonian mice, suggesting that PPX shifted choices towards immediate/small rewards at all delays (Fig. 1H). Interestingly, in sessions following a 48h PPX washout period, the *A* value recovered to baseline values, while differences in *K* and AUC persisted (Fig. S2H-J). These results suggest that in parkinsonian mice, PPX acutely reduces sensitivity to differences in reward magnitude, and chronically impairs sensitivity to delays. Together, these alterations may lead to impulsive decision making as seen in PD/ICD.

Clinical observations suggest vulnerability to ICD differs across individuals. Vulnerability has been associated with structural and functional deficits in brain regions related to reward processing, such as the caudate nucleus ^44^. To explore whether differences in baseline disease severity predicted vulnerability to ICD in our mouse model, we correlated postmortem measures of dopaminergic cell body and axonal integrity with key quantitative measures (*K & A*) associated with the delay discounting curve (Fig. S2K-O). A Spearman correlation analysis showed that in parkinsonian mice treated with PPX, *A* values were positively correlated with residual TH^+^ fluorescence in the DMS, but not in the DLS or VS (Fig. S2L), consistent with the idea that intact DMS dopaminergic signaling is crucial for encoding reward magnitudes ^45^. TH^+^ fluorescence in DMS or VS did not significantly correlate with *K* values (Fig. S2M). TH^+^ neurons in SNc did not significantly correlate with either *A* or *K* values (Fig. S2N,O). Altogether these findings demonstrate rodents can closely recapitulate key features of PD with ICD.

### Chemogenetic inhibition of iMSNs in the dorsomedial striatum mimics the effect of PPX in parkinsonian mice

ICD is reversible in people with PD upon dose reduction or discontinuation of dopamine agonist therapy ^19^, which implies ICD may be a disorder of drug-induced alterations in neural activity or connectivity. Though the specific brain areas or cell types which mediate ICD are unclear, neuroimaging and pharmacology provide some candidates. Clinically used dopamine agonists bind D2/3Rs ^46^. D2/3Rs are expressed across many brain regions including the striatum, amygdala, and hippocampus ^20,47,48^. While multiple brain areas have been implicated in ICD, several studies link alterations in striatal volume or functional connectivity to ICD ^49-51^. Within the striatum, D2Rs are most densely expressed on iMSNs of the indirect pathway ^20,52^. Within these neurons, dopamine signaling is hypothesized to reduce neural activity ^22-24^. However, the relationship of indirect pathway activity to ICD-related behavior remains unclear. To mimic the hypothesized effects of PPX on iMSN synaptic output, we used a chemogenetic (DREADD) approach. We expressed the inhibitory DREADD, hM4Di (Gi-coupled) or a control fluorophore (mCherry) in iMSNs of the DMS of parkinsonian mice (Fig. 2A). To validate the use of hM4Di, we first performed *ex vivo* whole-cell recordings from A2a-Cre;D2-eYFP mice coinjected with Cre-dependent ChR2-eYFP and Cre-dependent hM4Di (Fig. S3A), using the inhibitory connections between iMSNs and dMSNs as a functional readout of iMSN synaptic output. Brief light pulses evoked inhibitory postsynaptic currents (oIPSCs) in postsynaptic eYFP-negative dMSNs (Fig. S3B). Application of the DREADD agonist, clozapine-N-oxide (CNO), reduced oIPSC amplitude (Fig. S3B,C), confirming that the Gi-coupled DREADD inhibited iMSN output.

**Figure 2.**
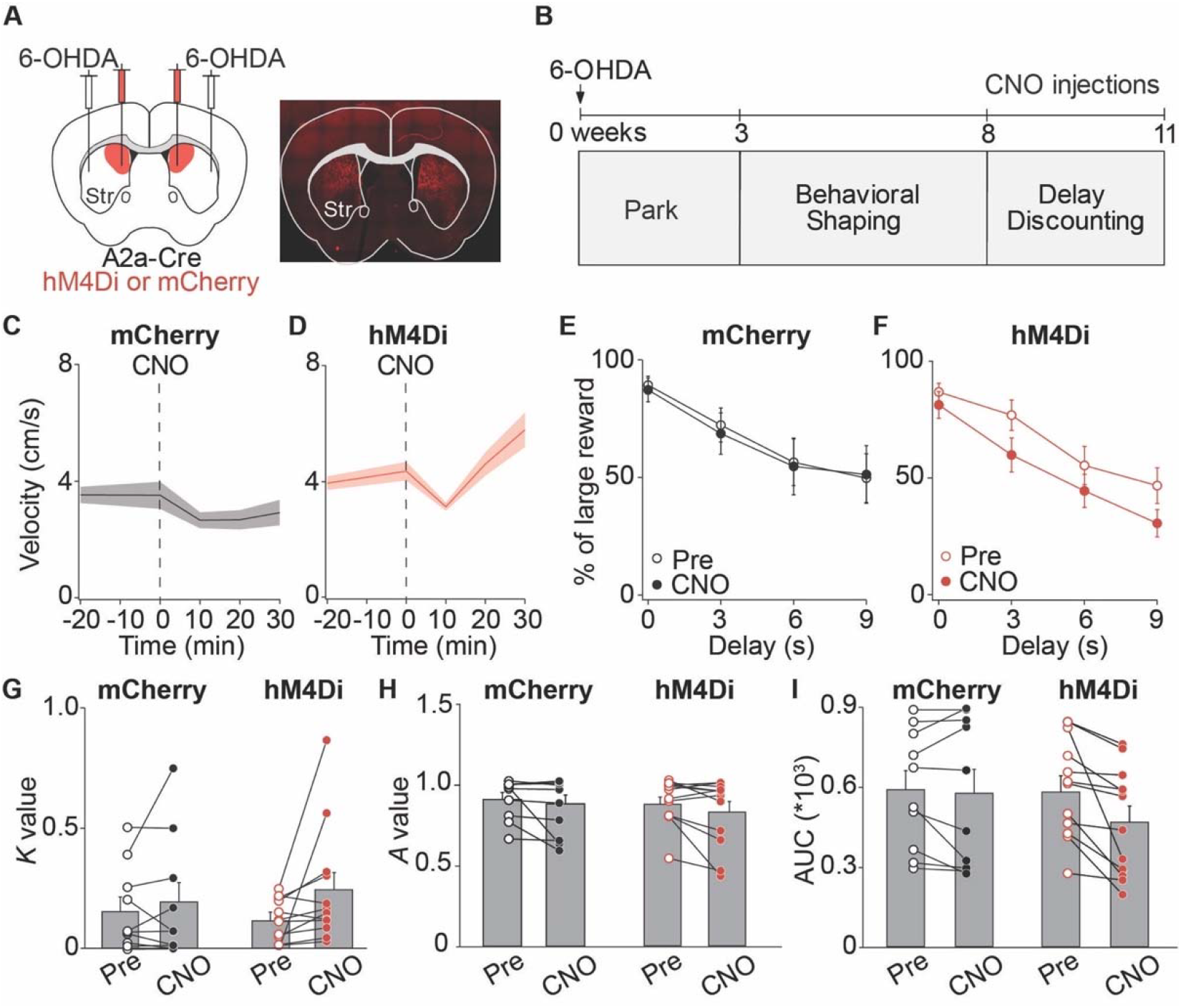
Chemogenetic inhibition of iMSNs in the dorsomedial striatum mimics the effect of PPX in parkinsonian mice. (**A**) Left: Injection schematic. Right: Postmortem tissue showing mCherry expression in the DMS (red). (**B**) Experimental timeline. (**C, D**) Locomotor activity following IP injection of CNO in parkinsonian mice expressing mCherry (C, N = 8, D, N = 13; baseline vs. post-CNO, p = 0.02). (**E, F**) Percentage delayed/large reward choices by either mCherry or hM4Di-expressing mice at each delay during Pre (open circles) and post-CNO injection (filled circles) sessions (E, N =10, p > 0.99 at all delays; F, N = 12, 0s: p = 0.61, 3s: p < 0.01, 6s: p = 0.11, 9s: p < 0.05). (**G-I**) *K, A* and AUC values from sessions before (Pre) and post-CNO administration in mice expressing mCherry or hM4Di (mCherry: N = 10, hM4Di: N = 12; G: mCherry, p = 0.63; hM4Di, p = 0.02; H: mCherry, p = 0.16; hM4Di, p = 0.10; I: mCherry, p = 0.43; hM4Di, p < 0.001). N, animals, all data presented as means ± SEMs.

We first tested whether chemogenetic inhibition of striatal iMSNs, like PPX, could ameliorate parkinsonian locomotor deficits. CNO treatment increased movement in the hM4Di, but not mCherry control group, suggesting a therapeutic effect (Fig. 2C,D). We then assessed whether chemogenetic inhibition of iMSNs is sufficient to cause impulsive decision-making. Parkinsonian hM4Di or mCherry mice were assessed in the delay discounting task, before and after CNO treatments. Chemogenetic inhibition of iMSNs robustly shifted choices towards immediate/small rewards over delayed/large rewards in hM4Di-expressing but not mCherry control mice (Fig. 2E,F). Moreover, chemogenetic inhibition of indirect pathway significantly increased the *K* value and decreased AUC, consistent with a greater degree of impulsivity (Fig. 2G,I). Interestingly, CNO did not impact *A* value in either group, suggesting that inhibition of the indirect pathway alone did not alter discrimination of reward sizes (Fig. 2H). Together, these findings suggest that chemogenetic inhibition of iMSNs within DMS is sufficient to induce impulsive decision-making in parkinsonian mice in the absence of PPX treatment.

### Pramipexole triggers bidirectional changes in striatal activity in parkinsonian mice

Similar to PPX, chemogenetic inhibition of striatal indirect pathway output increased impulsive decision-making. However, the responses of iMSNs (and dMSNs) to PPX in this model remain unknown. Dopamine D2/3R agonists like PPX may target D2Rs located on several microcircuit elements within the striatum ^53^. By acting on D2Rs, PPX may directly or indirectly influence the firing of dMSNs and iMSNs, which is important for appropriate decision-making ^54^. To determine how PPX affected dMSN and iMSN activity, we performed single-unit electrophysiology of optogenetically identified DMS neurons in both healthy and parkinsonian mice (Fig. 3A). Optogenetic labeling of dMSNs and iMSNs was achieved by expressing channelrhodopsin-2 (ChR2) selectively in dMSNs or iMSNs (using D1-Cre or A2a-Cre mice, respectively) ^55,56^ and recording light responses at the end of each session ^24,57^. We first determined whether dopamine loss caused changes in overall MSN activity, as the standard model predicts ^22^. We compared the firing rates of dMSNs and iMSNs in parkinsonian mice to those in healthy mice. The firing rates of DMS dMSNs and iMSNs in parkinsonian mice were very similar to those in control mice (Fig. 3B,C). These results indicate mild dopamine depletion does not markedly change overall MSN firing rates.

**Figure 3.**
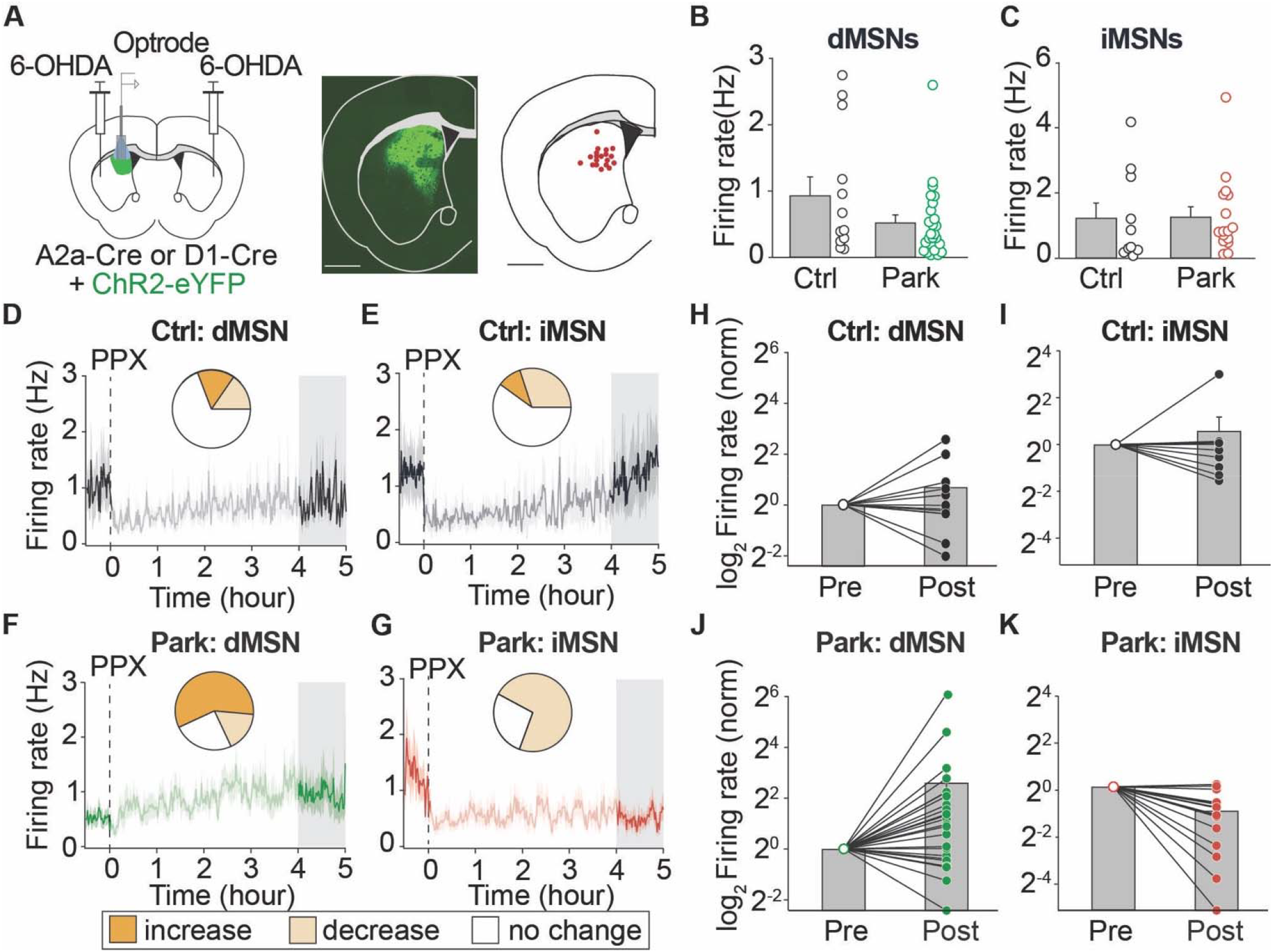
Pramipexole triggers bidirectional changes in striatal activity in parkinsonian mice. (**A**) Left: Schematic showing injection and optrode array implantation sites. Middle: Postmortem histology confirming the expression of ChR2-eYFP (green). Right: Recording sites verified by electrolytic lesions. (**B, C**) Average baseline firing rates of optogenetically labeled dMSNs (B) and iMSNs (C) in healthy and parkinsonian mice (B, Ctrl: [N = 3, n = 12] vs. Park: [N = 5, n = 24], p = 0.17; C, Ctrl: [N = 4, n = 10] vs. Park: [N = 4, n = 16], p = 0.70). (**D-G**) The effect of pramipexole (PPX) on optogenetically labeled dMSNs (D, F) and iMSNs (E, G). The shaded area at 4-5 hours post-injection represents the time of all behavioral experiments; firing rates were compared between baseline and this period (D: [N = 3, n = 12], p = 0.91; E: [N = 4, n = 10], p = 0.38; F: [N = 5, n = 24], p = 0.01; G: [N = 4, n = 16], p < 0.001). Insets: The proportion of optogenetically identified dMSNs and iMSNs whose firing rate increased, decreased, or had no response to PPX (D: increase: 15.4%, decrease: 15.4%, no change: 69.2%; E: increase: 10.0%, decrease: 30.0%, no change: 60.0%; F: increase: 58.3%, decrease: 16.7%, no change: 25.0%, p = 0.01; G: increase: 0%, decrease: 72.7%, no change: 27.3%, p = 0.002). (**H, I**) Summary of normalized response of dMSN (H) and iMSN (I) firing rates to PPX (compared to baseline) in healthy and parkinsonian mice (same data as displayed in D-G). N, animals, n, cells. All data presented as means ± SEMs.

To determine how PPX affected striatal firing over time, each recording session included a baseline period (30 min), PPX (0.5 mg/kg) injection, then a 5 h post-injection period. As delay discounting was tested between 4 and 5 h post-injection, we focused on the change between baseline and 4-5h (shaded area). In healthy mice, PPX suppressed activity in both dMSNs and iMSNs immediately after injection. However, at 4-5 h post-injection, the average firing rates of both cell types returned to baseline levels (Fig. 3D,E). We classified all optically-identified units into three categories based on PPX-induced changes in firing rate between baseline and 4-5 h post-injection: ‘increase’, ‘decrease’, and ‘no change’ (no significant difference) types. Responses in healthy mice were diverse: a small proportion of dMSNs were either inhibited or excited, but most dMSNs showed no change in firing rate. iMSNs showed similar variability (Fig. 3D,E, insets). In parkinsonian mice, however, PPX caused bidirectional changes in optically labeled MSN firing rates. PPX increased dMSN firing rates and decreased iMSN firing rates (Fig. 3F,G). Moreover, both dMSNs and iMSN from parkinsonian mice exhibited more pronounced changes in response to PPX than in healthy mice (Fig. 3H-K). This phenomenon is reflected in the larger percentage of ‘increase’ type dMSNs and ‘decrease” type iMSNs in parkinsonian compared to healthy mice (Fig. 3F, G insets).

Importantly, similar patterns were seen in the larger unlabeled MSN pool to those in the smaller optogenetically labeled pool (Fig. S4A-D). Given the potential variability in firing rates over prolonged recordings, in separate experiments we injected saline instead of PPX. Nearly all units showed no change in firing rates (Fig. S4E,F). These findings demonstrate that MSNs are indeed bidirectionally dysregulated by PPX in parkinsonian mice, indicating aberrant MSN activity is a potential driver for ICD.

### Impulsive decision-making develops over successive doses of pramipexole, in parallel with changes in striatal activity

Chronic dopamine replacement therapies (levodopa and dopamine agonists), may cause involuntary movements and cognitive-behavioral dysfunction, highlighting plasticity at the behavioral level ^58,59^. At the cellular level, MSNs are highly dependent on the surrounding local circuitry to drive spiking activity ^60^. Striatal circuits are known to be regulated by dopamine on both acute and chronic timescales ^23,53,61,62^. Indeed, alterations in receptor expression, corticostriatal input, and local inhibitory connections have been reported in dopamine depleted animals undergoing dopamine replacement therapy ^63-65^. These forms of plasticity may cause an augmented response to dopamine over successive exposures, potentiating abnormal behaviors. In fact, chronic treatment with dopamine agonists result in more profound changes in behavior ^8,66,67^. To determine whether chronic PPX treatment exacerbated changes in delay discounting, we compared behavior across four PPX injection sessions (Fig. 4A). In healthy control mice, delay discounting remained consistent across sessions (Fig. 4B,S5A). However, in parkinsonian mice, delay discounting changed over time: PPX injection induced a modest increase in impulsivity, which became more marked by the 4^th^ session (Fig. 4C,S5B). Repeated PPX treatment resulted in larger changes in delay discounting behavior in parkinsonian mice.

**Figure 4.**
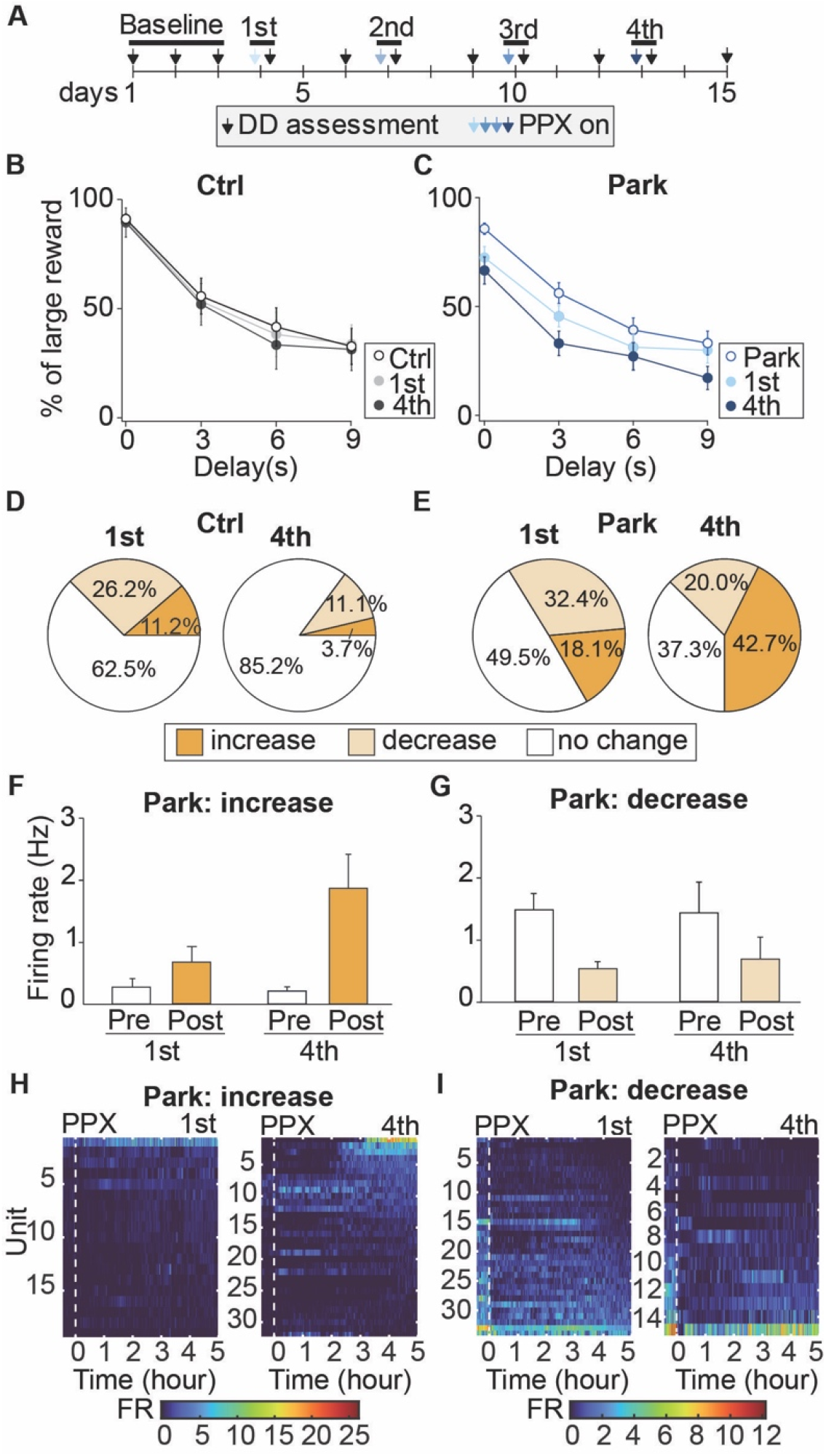
Impulsive decision-making develops over successive doses of pramipexole, in parallel with changes in striatal activity. (A-C) In healthy control and parkinsonian mice, delay discounting behavior was measured in PPX-naïve mice (baseline) and across 4 PPX treatment sessions. (A) Experimental timeline. (B-C) Percentage of trials in which healthy control (B) and parkinsonian (C) mice chose the delayed/large reward across delays during baseline (open circles) and in the1^st^ and 4^th^ PPX session (filled circles) (Ctrl: N = 16, Park: N = 31; B, 1^st^ vs. 4^th^ : p > 0.99 at all delays; C, 1^st^ vs. 4^th^ : 0s: p = 0.47, 3s: p < 0.05, 6s: p > 0.99, 9s: p < 0.05, for other comparisons, refer to statistical table). (D-E) Proportion of each response type during after the 1^st^ and 4^th^ PPX session in control and parkinsonian mice (Ctrl: 1^st^ [N = 7, n = 80] vs. 4^th^ [N = 4, n = 27], p = 0.11; Park: 1^st^ [N = 10, n = 105] vs. 4^th^ [N = 8, n = 75], p = 0.002). (F-G) Average firing rates before and after PPX among each response type in parkinsonian mice (F: 1^st^ [N = 6, n = 19] vs. 4^th^ [N = 7, n = 32], p = 0.02; G: 1^st^ [N = 9, n = 34] vs. 4^th^ [N = 5, n = 15], p = 0.89). (H-I) Heatmaps showing firing rates over time following PPX injection in parkinsonian mice. Responses during the 1^st^ PPX session are at left, the 4^th^ session at right, for neurons with an increase (H) or decrease (I) type response. Each row represents a single unit. N, animals, n, cells. All data presented as means ± SEMs.

To determine whether plasticity in the responses of MSNs to PPX might underlie this behavioral plasticity, we compared how MSN firing changed between the 1^st^ and 4^th^ PPX session. In control mice, the proportion of response types was consistent across injection days (Fig. 4D,S6A). However, in parkinsonian mice, the response types shifted over four sessions (Fig. 4E,S6B). Notably, ‘increase’ type MSNs showed more dramatic increases in firing rate in response to the 4^th^ (versus 1^st^) PPX injection (Fig. 4F,S6C), while ‘decrease’ type MSNs responded similarly across sessions (Fig. 4G,S6D). Together, these findings indicate chronic PPX treatment leads to a higher proportion of excited MSNs, each of which has a more dramatic response (Fig. 4H,S6E). Conversely, the proportion of ‘decrease’ MSNs falls over PPX treatment (Fig. 4I,S6F). Together, these findings indicate that in parkinsonian animals, dopamine agonists lead to changes in striatal firing which may contribute to the development of ICD-like behavior.

## DISCUSSION

Here, we established a mouse model of impulse control disorder (ICD) in Parkinson’s disease (PD) and investigated the role of aberrant striatal activity in impulsive decision-making. We found that pramipexole (PPX), a widely used D2/3R agonist associated with high risk of ICD ^4,5^, induced impulsive decision-making in parkinsonian mice, as assessed by the delay discounting task. Inhibition of indirect pathway output was sufficient to cause impulsive decision-making, mimicking the effects of PPX. PPX caused bidirectional changes in iMSN/dMSN firing rates in parkinsonian mice, while having minimal effects in healthy mice. Chronic PPX treatment potentiated changes in striatal physiology and decision-making behavior. Our study is the first to perform physiological recordings in a clinically-relevant mouse model of ICD, and to link a specific striatal pathway to ICD.

We found that in mildly parkinsonian mice, dopamine agonist treatment reproduced key clinical features of ICD. Significant motor deficits can create confounds in operant tasks, preventing accurate assessment of cognitive-behavioral phenotypes in mouse models of PD. To overcome this potential obstacle, we adapted a mouse model of PD with relatively restricted bilateral dopamine depletion, reminiscent of what is seen in early PD, when dopamine agonists are most likely to be employed. While this model showed mild motor deficits, animals could still learn and perform our task, and motivational metrics were comparable to those in control mice. Our model reflected key clinical features of PD/ICD, such as the increased risk in PD patients (versus healthy individuals) and the medication dependence of impulsive behavior ^10-12^. A key variable in our model was the dose of dopamine agonist. Prior work indicates higher doses D2/3R agonist have reinforcing properties even in intact animals ^68,69^. We calibrated the agonist to provide motor benefit while avoiding supratherapeutic dosing. Overall, we believe that the risk of ICD is associated with increased dopamine signaling in a vulnerable neural substrate, which would explain the differences between healthy and parkinsonian mice in their behavioral and physiological responses to PPX.

We used delay discounting behavior as an assay of impulsive decision-making, as changes in delay discounting have been observed in people with PD/ICD ^10-12^. However, this assay reflects only one facet of ICD. ICD-related behaviors encompass motor impulsivity (impulsive actions) and decision impulsivity (impulsive choices) ^70,71^. Other tasks that might be used to capture other features of ICD include the 5-choice serial reaction time task, which assesses motor impulsivity ^8^, the probability discounting task, which evaluates risky choice ^66^, or rodent versions of the Iowa Gambling Task, which is abnormal in PD/ICD ^72^, and can mimic the salient sensory stimuli and rewards of a casino ^73^.

Altered delay discounting behavior can be driven by changes in how reward magnitude, time, and/or reward/delay tradeoffs are processed ^74^. As in previous studies, we used Herrnstein’s hyperbolic model V = *A*/(1+*K*D to fit behavior ^33,34,39^. In this equation, ‘D’ represents the delay, and ‘*A*’ and ‘*K*’ factors reflect how mice perceive different reward sizes and delay durations, respectively. We found that PPX increased *K* value in parkinsonian mice, much as has been seen in PD patients with ICD ^6,11,12^. These findings suggest PPX-treated mice are more intolerant of waiting, even for a larger reward. Impulsivity also correlates with poor temporal discrimination in rats and humans ^74,75^. Interestingly, in a subset of parkinsonian mice, PPX also decreased *A* value, suggesting impaired processing of reward magnitude. While the small number of delay discounting studies in PD/ICD have not shown changes in the *A* value, other studies of impulsivity suggest reward magnitude discrimination is crucial in driving impulsive choice ^41^. We suspect this difference may be related to differences in human versus rodent delay discounting tasks or to the pattern of striatal dopamine depletion. We found that animals with greater dopaminergic denervation in the dorsomedial striatum (DMS) tended to have changes in *A* value in PPX-treated parkinsonian mice. These findings are in line with studies that indicate the DMS encodes reward magnitudes ^45^.

We found that chemogenetic inhibition of DMS iMSNs induced an ICD-like phenotype of more pronounced delay discounting. This observation is in line with evidence that the associative striatum (caudate nucleus in primates, or DMS in rodents) plays a significant role in mediating impulsive decision-making, including pharmacological and electrophysiological studies linking this region to delay discounting and decision-making in healthy animals ^13,15,76^. It is also consistent with the pharmacology of PPX and iMSNs. PPX would be predicted to reduce indirect pathway output, while disinhibiting direct pathway activity via local inhibitory collaterals ^77^. Prior work has demonstrated that D2/3R agonists increase activity in the globus pallidus (GP), and decrease activity in the substantia nigra reticulata (SNr) in monkeys ^78,79^. Chemogenetic inhibition of iMSNs may mimic some effects of PPX, leading to impulsive decision-making. However, there were differences in the behavioral effects of chemogenetic manipulation of iMSNs and PPX. In parkinsonian mice, PPX decreased *A* and increased *K* values; chemogenetic inhibition only increased *K* values. This discrepancy may be explained by the effects of PPX outside iMSNs and/or the striatum, including through D2Rs on frontal cortical neurons (and their terminals in the striatum) critical for decision-making ^52,80,81^. Alternatively, differences may arise from the action of PPX on D3Rs, which colocalize with D1Rs in the ventral striatum ^82^, but whose expression is increased in the dorsal striatum in parkinsonian animals treated with dopamine replacement therapy ^65^.

We found that the dopamine agonist PPX induced changes in striatal activity in parkinsonian mice, providing a potential substrate for ICD-like behavior. This dysregulation is likely to arise from the interaction of PPX with the chronically dopamine-depleted striatum. Previous work has identified many alterations to striatal signaling molecules and physiological properties in people with PD and animal models of PD ^83,84^. These alterations include upregulation of D2Rs ^85,86^, which may explain the more pronounced suppression of iMSN firing in parkinsonian mice. dMSN sensitivity to PPX could be mediated by suppression of collateral inhibition on both acute and chronic timescales ^87^. In healthy mice, striatal activity encodes key aspects of delay discounting behavior, including reward size and elapsed waiting time ^16,18,45^. PPX disrupted striatal activity in our mouse model of PD/ICD, which may enhance the perceived value of immediate rewards and/or impair learning from waiting time, biasing animals towards immediate/small rewards. Future studies of striatal activity during behavioral tasks in the PD/ICD model may reveal the precise mechanisms by which PPX alters decision-making. We also found that over multiple doses, behavioral and physiological responses to PPX potentiated. This may relate to additional adaptations in striatal circuitry, as have been seen with repeated dopaminergic treatments in animal models of psychostimulant sensitization, chronic PPX treatment ^88-90^, or levodopa-induced dyskinesia ^65^.

Together, our results suggest a key potential mechanism for impulsive decision-making in ICD: dysregulated dMSN and iMSN activity in parkinsonian animals treated with dopamine agonist medication. This insight could inform the use of dopamine replacement therapy with a goal of preventing or ameliorating ICD.

## Data Availability

Datasets are available at https://doi.org/10.5281/zenodo.10703094. Detailed protocols and analysis code are listed within ‘METHOD DETAILS’ section.

## ACKNOWLEDGEMENTS

The authors wish to thank Drs. Elyssa Margolis, Joshua Berke, and Vijay Mohan K Namboodiri, as well as members of the Nelson laboratory, for their valuable feedback on this manuscript. This work was supported by grants from the Parkinson’s Foundation (PF-PRF-836910 to X.Z.), the National Institute of Neurological Disorders and Stroke (NINDS) (R01NS101354 to A.B.N.), Aligning Science Across Parkinson’s (ASAP-020529 to A.B.N.) through the Michael J. Fox Foundation for Parkinson’s Research (MJFF), and by the Richard and Shirley Cahill Endowed Chair in Parkinson’s Disease Research (to A.B.N.).

