## Supplemental Figures and Tables for "Inhibition of Indirect Pathway Activity Causes Abnormal Decision-Making In a Mouse Model of Impulse Control Disorder in Parkinson’s Disease"

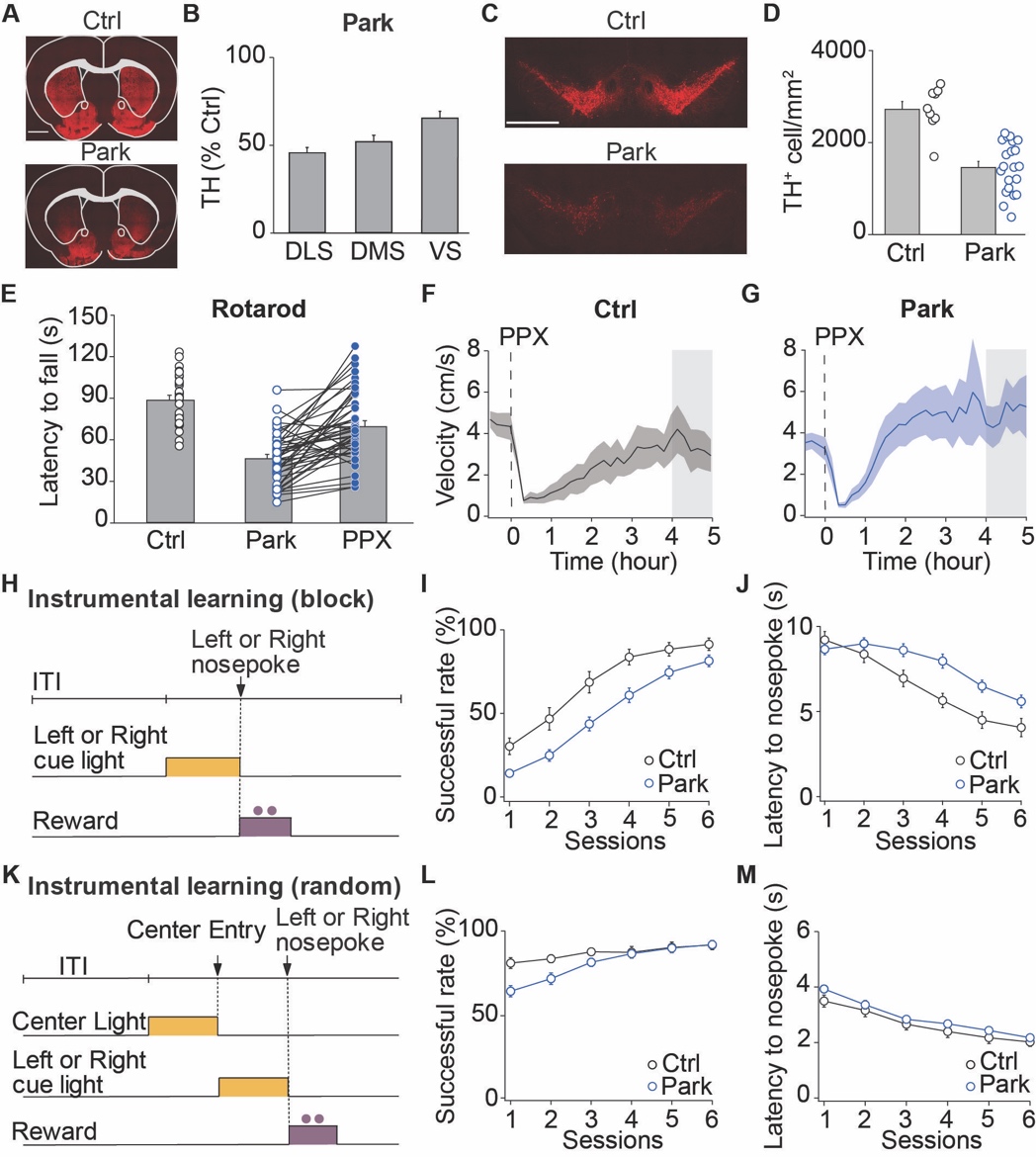

**Figure S1 (Associated with Figure 1).**

(**A-D**) Postmortem coronal sections containing the striatum (A) and midbrain (C) were immunostained for tyrosine hydroxylase (TH, red). (**A**) Representative striatal sections from control (top) and parkinsonian (bottom) mice. (**B**) Average TH^+^ immunofluorescence across different striatal subregions in parkinsonian mice, normalized to saline-injected control sections (N = 28, DMS vs. DLS: p = 0.56; VS vs. DMS: p < 0.03; VS vs. DLS: p < 0.001). (**C**) Representative sections containing the substantia nigra pars compacta (SNc) from control (top) and parkinsonian (bottom) mice. (**D**) Density of TH^+^ dopaminergic neurons within the SNc, normalized to the measured area (Ctrl: N = 9, Park: N = 21, p < 0.001). (**E**) On the accelerating rotarod test, parkinsonian mice showed a lower latency to fall, but improved with PPX (0.5 mg/kg) (Ctrl: N = 31, Park: N = 45, PPX: N = 45; Ctrl vs. Park: p < 0.001, Park vs. PPX: p < 0.001). (**F, G**). Open field locomotor activity in control (F) and parkinsonian (G) mice following PPX injection. Operant behavior was tested during the shaded period (4-5 hours post-PPX) (F, N = 7; G, N = 13). (**H-M**) Detailed metrics from behavioral shaping (instrumental learning). (**H**) The instrumental learning (blocked) task structure. (**I**) The success rate (cued side nosepoke) increased in both healthy and parkinsonian mice across instrumental learning sessions (Ctrl: N = 24, Park: N = 51; session 1, 3&4: p < 0.05, session 2, 5&6: p > 0.05). (**J**) Average time between side cue light illumination and side nosepoke in instrumental learning sessions (session 1-2 & 6, p > 0.05; session 3-5, p < 0.05). (**K**) The instrumental learning (random) task structure. (**L**) The success rate (trial self-initiation followed by cued side nosepoke) was high across instrumental learning sessions (Ctrl: N = 24, Park: N = 51; session 1: p < 0.05, session 2-6: p > 0.05). (**M**) Average time between central cue light illumination and trial initiation (session 1: p < 0.05, session 2-6: p > 0.05). N, animals, all data presented as means ± SEMs. Scale bars, 1mm.

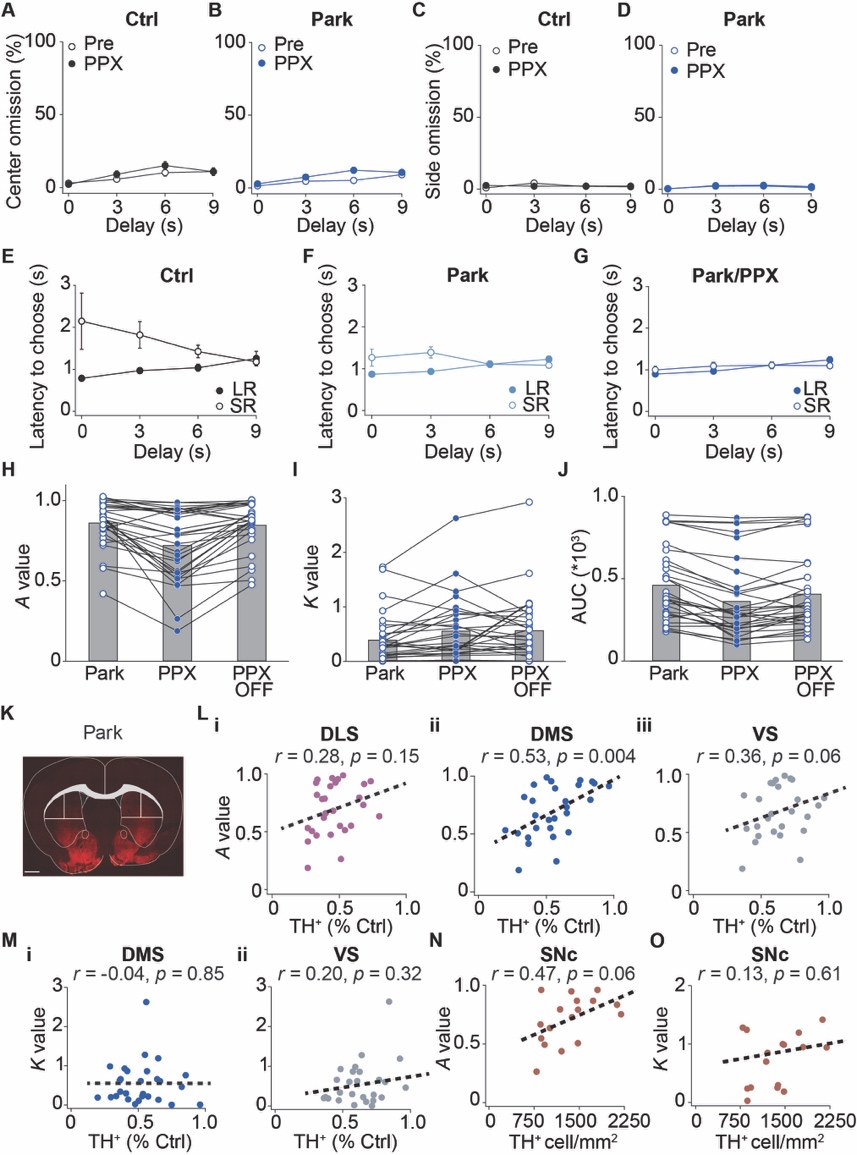

**Figure S2 (Associated with Figure 1). Detailed metrics from the delayed discounting task.**

(**A, B**) Percentage of free choice trials in which mice did not initiate the trial (Ctrl: N = 16, Park: N = 31; A, p > 0.05 at all delays; B, p > 0.05, except 6s: p < 0.05). (**C, D**) Percentage of free choice trials in which mice did not make a side (choice) nosepoke Ctrl: N = 16, Park: N = 31; C, p > 0.05 at all delays; D, p > 0.05 at all delays). (**E-G**) Time between cue lights and choice nosepoke for delayed/large rewards (filled circles) and immediate/small rewards (open circles) at each delay during free choice trials in healthy control (E), parkinsonian (F) and PPX-treated parkinsonian mice (G) (Ctrl: N = 16, Park: N = 31; LR vs. SR, E, 0s: p = 0.12, 3s: p < 0.01, 6s: p < 0.05, 9s: p > 0.99; F, 0s: p = 0.77, 3s: p < 0.05, 6s: p > 0.99, 9s: p = 0.56; G, p > 0.99 at all delays). (**H-J**) *A*, *K* and AUC values in all three conditions in parkinsonian mice: baseline, post-PPX treatment, and PPX washout (48 hours) (N = 31, Park vs. PPX off, H: p > 0.99; I: p = 0.01; J: p < 0.01). (**K**). A representative coronal section from a parkinsonian mouse, immunolabeled for TH, illustrating the delineation of subregions within the striatum: dorsomedial striatum (DMS), dorsolateral striatum (DLS), and ventral striatum (VS). (**Li-iii**). Scatter plots and fitted line demonstrating the correlation between the *A* values on PPX and residual TH^+^ fluorescence intensity in the subregions of the striatum in parkinsonian mice (N = 28; i: p = 0.15, r = 0.28, r^2^ = 0.08; ii: p = 0.004, r = 0.53, r^2^ = 0.28; iii: p = 0.06, r = 0.36, r^2^ = 0.13). (**Mi-ii**). Scatter plots and fitted line demonstrating the correlation between the *K* values on PPX and residual TH^+^ fluorescence intensity in the subregions of striatum in parkinsonian mice (N = 28; i: p = 0.85, r = -0.04, r^2^ = 0.001; ii: p = 0.32, r = 0.20, r^2^ = 0.04). (**N**). Scatter plot and fitted line demonstrating the correlation between *A* values on PPX and SNc dopaminergic neuron density of parkinsonian mice (N = 17; p = 0.06, r = 0.47, r^2^ = 0.22). (**O**). Scatter plot and fitted line demonstrating the correlation between *K* values on PPX and SNc dopaminergic neurons density of parkinsonian mice (N = 17; p = 0.61, r = 0.13, r^2^ = 0.02). N, animals, all data presented as means ± SEMs.

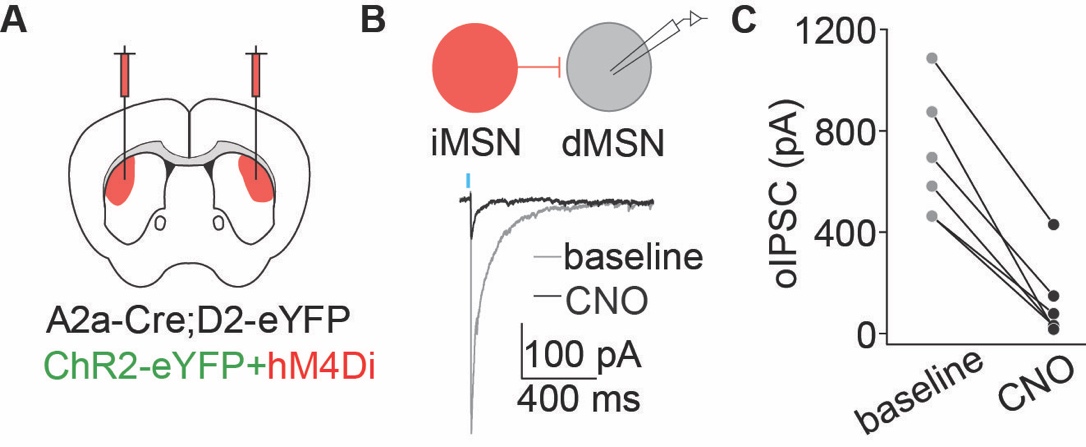

**Figure S3 (Associated with Figure 2).**

(**A**) Schematic diagram showing A2a-Cre;D2-eYFP mice that were bilaterally injected with DIO-hM4D(Gi) into the DLS for validating the hM4Di virus via slice electrophysiology. (**B**) Top: schematic diagram showing identification of eYFP-negative dMSNs and the recording configuration. Bottom: representative electrophysiological trace showing oIPSC before (black) and after CNO (grey, 1 μM) bath-application.(**C**) Summary oIPSC amplitudes before and after CNO bath-application (N = 2, n = 6, p < 0.01). N, animals, n, cells. All data presented as means ± SEMs. This slice validation data was also used in another manuscript^31^.

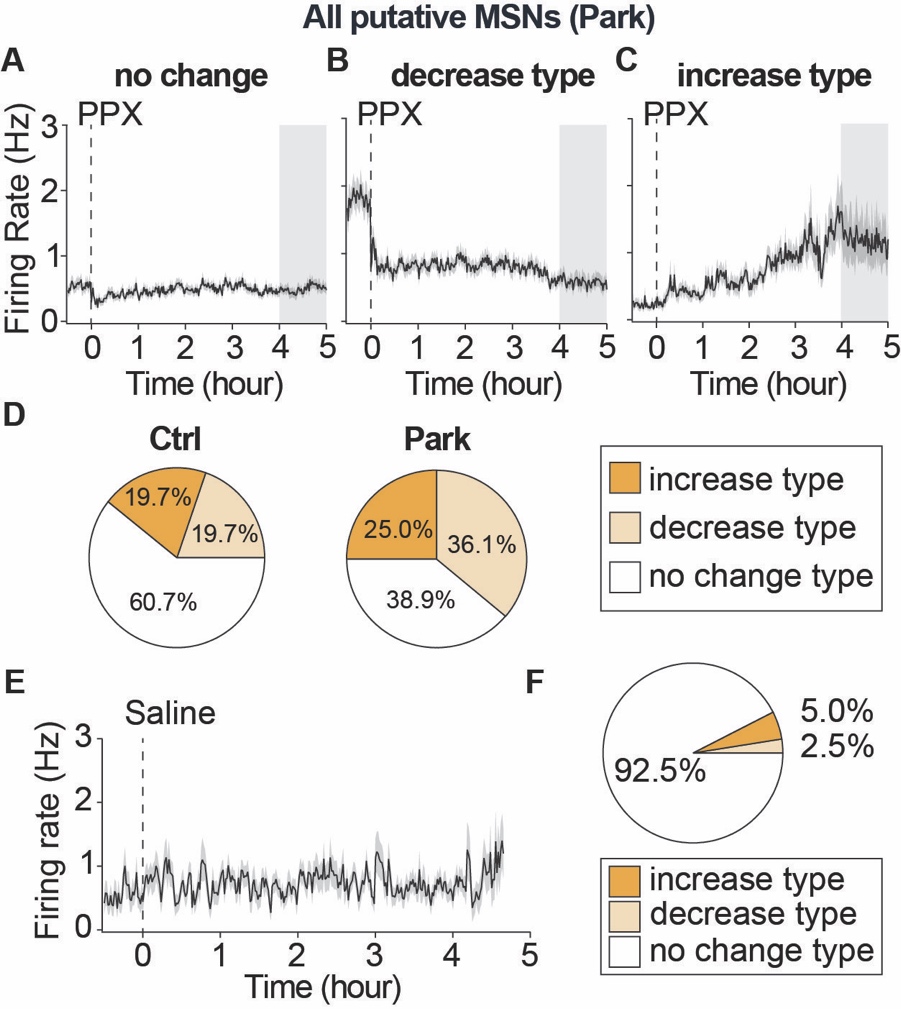

**Figure S4 (Associated with Figure 3)**

(**A-C**). Average firing rates of putative MSNs from parkinsonian mice, categorized based on response type as 'increase' (A), 'decrease' (B), or 'no change' (C) (N = 11, increase: n = 54, decrease: n = 78, no change: n = 82). (**D**). Proportion of putative MSNs classified by changes in firing rate in response to PPX (Ctrl, [N = 7, n = 117] vs. Park, [N = 11, n = 214], p = 0.0003). (**E**). Average firing rate of putative MSNs from parkinsonian mice over time in response to saline administration (dotted line: saline i.p. injection, [N = 4, n = 40]). (**F**). Proportion of putative MSNs categorized by changes in firing rate following saline administration (increase: n = 1, decrease: n = 2, no change: n = 37), N, animals, n, cells. All data presented as means ± SEMs. This slice validation data is also used in another manuscript ^31^.

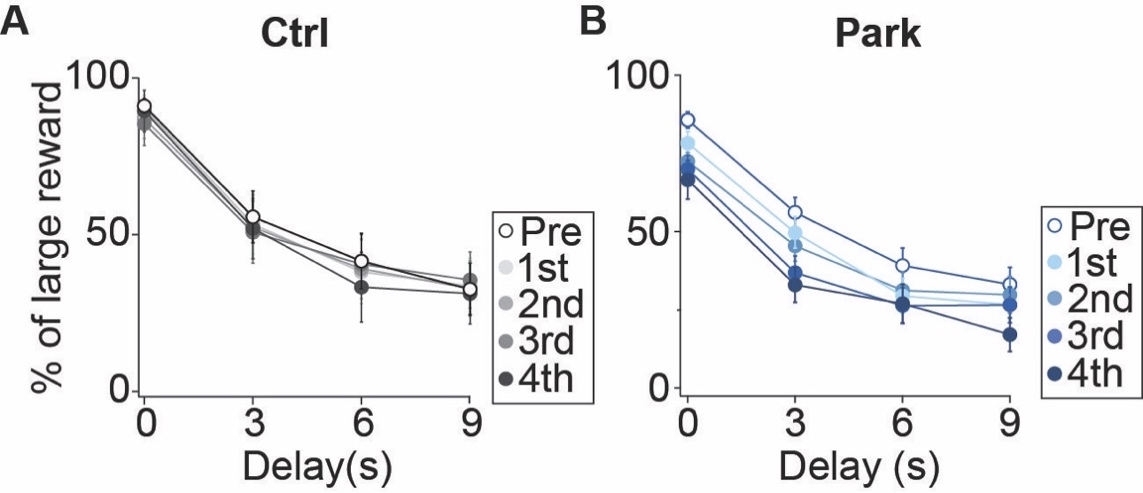

**Figure S5 (Associated with Figure 4). (**A-B) Percentage of trials in which healthy control (A) and parkinsonian (B) mice chose the delayed/large reward across delays during baseline (open circles) and the 1^st^ through 4^th^ PPX sessions. N, animals, all data presented as means ± SEMs.

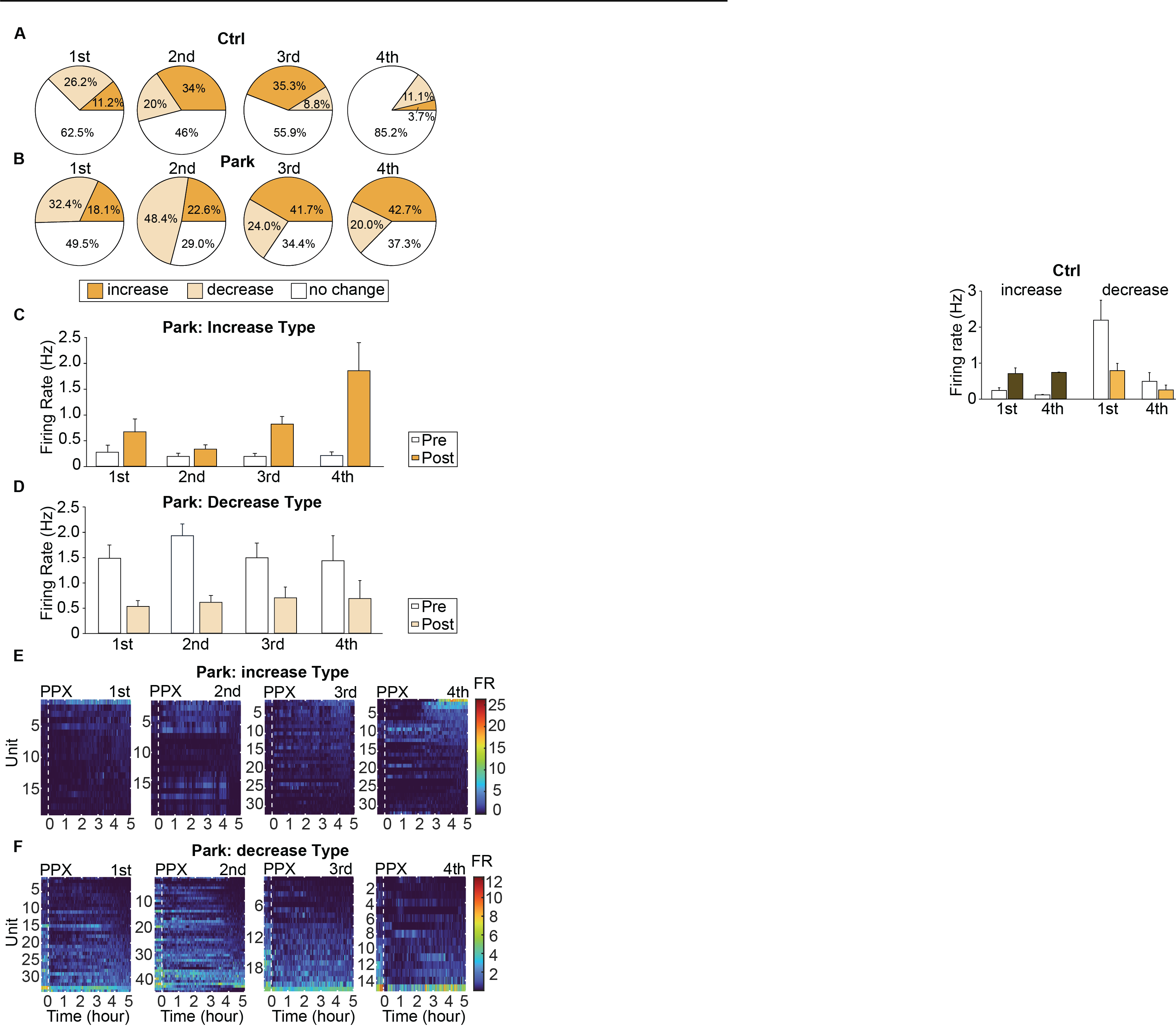

**Figure S6 (Associated with Figure 4).** (A-B) Proportion of putative MSNs with an increase, decrease, or no change in firing rate in response to PPX (assessed 4 hours post-injection) in control (A) and parkinsonian (B) mice. (C-D) Average firing rates of 'increase' (C) or 'decrease' type (D) MSNs pre-PPX and across four post-PPX epochs in parkinsonian mice. (E-F). Heatmaps showing firing rates over time during 4 PPX injection sessions. Responses during the four sessions, are from left to right, for neurons with an increase (E) or decrease (F) type response. Each row represents a single unit. N, animals, n, cells. All data presented as means ± SEMs.

| **Table 1. EXPERIMENTAL ANALYSIS AND STATISTICS**  WSR, Wilcoxon Signed Rank test; MWU; Wilcoxon Rank Sum or Mann–Whitney U test; KW, Kruskal-Wallis test | | | | | | |
| --- | --- | --- | --- | --- | --- | --- |
| **Key Experiments** | **Figure** | **Comparison** | **Statistical**  **test** | **N (animals)** | **n (units)** | ***p* value** |
| PPX effects on DD curve  (Ctrl) | Fig. 1C | Within animal | WSR, Bonferroni | N = 16 | NA | 0s: p > 0.99  3s: p > 0.99  6s: p > 0.99  9s: p > 0.99 |
| PPX effects on DD curve  (Park) | Fig.1D | Within animal | WSR, Bonferroni | N = 31 | NA | 0s: p < 0.001  3s: p < 0.001  6s: p < 0.001  9s: p < 0.01 |
| PPX effects on *A* value  (Park) | Fig. 1F | Within animal | WSR, | N = 31 | NA | P < 0.001 |
| PPX effects on *K* value  (Park) | Fig. 1G | Within animal | WSR, | N = 31 | NA | P = 0.01 |
| PPX effects on AUC value  (Park) | Fig. 1H | Within animal | WSR, | N = 31 | NA | P < 0.001 |
| TH^+^ fluorescence | Fig. S1B | Between group | KW  Dunn's | N = 28 | NA | DMS vs. DLS: p = 0.56  DMS vs. VS: p <0.03  DLS vs. VS: p < 0.001 |
| SNc cell density  (Ctrl vs. Park) | Fig. S1D | Between group | MWU | Ctrl: N = 9  Park: N = 21 | NA | P < 0.001 |
| PPX effects on rotarod | Fig. S1E | Between group | KW  Dunn's | Ctrl: N = 31  Park: N = 45  PPX: N = 45 | NA | Ctrl vs. Park: p < 0.001  Park vs. PPX: p < 0.001  Ctrl vs. PPX: p < 0.01 |
| Instrumental  learning (block)  (Ctrl vs. Park) | Fig. S1I | Between group | MWU,  Bonferroni | Ctrl:  N = 24  Park:  N = 51 | NA | session 1: p < 0.01  session 2: p = 0.06  session 3: p < 0.05  session 4: p < 0.01  session 5: p = 0.14  session 6: p = 0.11 |
| Instrumental learning (block)  Latency to nosepoke  (Ctrl vs. Park) | Fig. S1J | Between group | MWU,  Bonferroni | Ctrl:  N = 24  Park:  N = 51 | NA | session 1: p >0.99  session 2: p > 0.99  session 3: p < 0.05  session 4: p < 0.01  session 5: p < 0.05  session 6: p = 0.10 |
| Instrumental learning (random)  (Ctrl vs. Park) | Fig. S1L | Between group | MWU,  Bonferroni | Ctrl:  N = 24  Park:  N = 51 | NA | session 1: p = 0.02  session 2: p = 0.68  session 3: p > 0.99  session 4: p > 0.99  session 5: p > 0.99  session 6: p > 0.99 |
| Instrumental learning  (random)  reaction time  (Ctrl vs. Park) | Fig.S1M | Between group | MWU,  Bonferroni | Ctrl:  N = 24  Park:  N = 51 | NA | session 1: p < 0.05  session 2: p > 0.99  session 3: p > 0.99  session 4: p > 0.99  session 5: p = 0.75  session 6: p > 0.99 |
| PPX effects on center omission (Ctrl) | Fig. S2A | Within animal | WSR, Bonferroni | N = 16 | NA | 0s: p > 0.99  3s: p > 0.99  6s: p = 0.70  9s: p > 0.99 |
| PPX effects on center omission(Park) | Fig. S2B | Within animal | WSR, Bonferroni | N = 31 | NA | 0s: p = 0.15  3s: p = 0.21  6s: p < 0.01  9s: p > 0.99 |
| PPX effects on side omission (Ctrl) | Fig. S2C | Within animal | WSR, Bonferroni | N = 16 | NA | 0s: p = 0.88  3s: p = 0.99  6s: p > 0.99  9s: p > 0.99 |
| PPX effects on side omission (Park) | Fig. S2D | Within animal | WSR, Bonferroni | N = 31 | NA | 0s: p > 0.88  3s: p > 0.99  6s: p > 0.99  9s: p = 0.92 |
| PPX effects on latency to trigger  (Ctrl) | Fig. S2E | Within animal | WSR, Bonferroni | N = 16 | NA | 0s: p = 0.12  3s: p < 0.01  6s: p < 0.05  9s: p >0.99 |
| PPX effects on latency to trigger  (Park ) | Fig. S2F | Within animal | WSR, Bonferroni | N = 31 | NA | 0s: p = 0.77  3s: p < 0.05  6s: p >0.99  9s: p = 0.56 |
| PPX effects on latency to trigger  (Park + PPX) | Fig.S2G | Within animal | WSR, Bonferroni | N = 31 | NA | 0s: p >0.99  3s: p >0.99  6s: p >0.99  9s: p >0.99 |
| *A* value during PPX off  (Park) | Fig. S2H | Within animal | WSR  Bonferroni | N = 31 | NA | PPX vs. PPX off: p < 0.001  Park vs. PPX off: p >0.99 |
| *K* value during PPX off  (Park) | Fig. S2I | Within animal | WSR  Bonferroni | N = 31 | NA | PPX vs. PPX off: p > 0.99  Park vs. PPX off: p = 0.01 |
| *AUC* value during PPX off  (Park) | Fig. S2J | Within animal | WSR Bonferroni | N = 31 | NA | PPX vs. PPX off: p < 0.01  Park vs. PPX off: p < 0.01 |
| Correlation: *A* and TH^+^ DLS | Fig.S2Li | NA | Pearson’s | N = 28 | NA | r = 0.28  p = 0.15  r^2^ = 0.08 |
| Correlation: *A* and TH^+^ DMS | Fig. S2Lii | NA | Pearson’s | N = 28 | NA | r = 0.53  p = 0.004  r^2^ = 0.28 |
| Correlation: *A* and TH^+^ VS | Fig. S2Liii | NA | Pearson’s | N = 28 | NA | r = 0.36  p = 0.06  r^2^ = 0.13 |
| Correlation: *K* and TH^+^ DMS | Fig. S2Mi | NA | Pearson’s | N = 28 | NA | r = -0.04  p = 0.85  r^2^ = 0.001 |
| Correlation: *K* and TH^+^ VS | Fig. S2Mii | NA | Pearson’s | N = 28 | NA | r = 0.20  p = 0.32  r^2^ = 0.04 |
| Correlation: *A* and TH^+^ SNc | Fig. S2N | NA | Pearson’s | N = 17 | NA | r = 0.47  p = 0.06  r^2^ = 0.22 |
| Correlation: *K* and TH+ SNc | Fig. S2O | NA | Pearson’s | N = 17 | NA | r = 0.13  p = 0.61  r^2^ = 0.02 |
| Locomotion  (hM4Di) | Fig. 2D | Within animal | WSR | N = 13 | NA | p = 0.02 |
| CNO effects on DD curve  (mCherry) | Fig. 2E | Within animal | WSR, Bonferroni | N = 10 | NA | 0s: p >0.99  3s: p >0.99  6s: p >0.99  9s: p >0.99 |
| CNO effects on DD curve  (hM4Di) | Fig. 2F | Within animal | WSR, Bonferroni | N = 12 | NA | 0s: p = 0.61  3s: p < 0.01  6s: p = 0.11  9s: p < 0.05 |
| CNO effects on *K* value  (mCherry) | Fig. 2G | Within animal | WSR | N = 10 | NA | p = 0.63 |
| CNO effects on *K* value  (hM4Di) | Fig. 2G | Within animal | WSR | N = 12 | NA | p = 0.02 |
| CNO effects on *A*  (mCherry) | Fig. 2H | Within animal | WSR | N = 10 | NA | p = 0.16 |
| CNO effects on *A*  (hM4Di) | Fig. 2H | Within animal | WSR | N = 12 | NA | p = 0.10 |
| CNO effects on AUC  (mCherry) | Fig. 2I | Within animal | WSR | N = 10 | NA | p = 0.43 |
| CNO effects on AUC  (hM4Di) | Fig. 2I | Within animal | WSR | N = 12 | NA | p < 0.001 |
| Inhibitory DREADD on oIPSCs | Fig. S3C | Within cell | WSR | N = 2 | n = 6 | p < 0.01 |
| dMSNs FR:  Ctrl vs. Park | Fig. 3B | Between group | MWU | Ctrl: N = 3  Park: N = 5 | Ctrl:  n = 12  Park:  n = 24 | p = 0.17 |
| iMSNs FR:  Ctrl vs. Park | Fig. 3C | Between group | MWU | Ctrl: N = 4  Park: N = 4 | Ctrl:  n = 10  Park:  n = 16 | p = 0.70 |
| PPX effects on dMSNs (Ctrl) | Fig. 3D | Within unit | WSR | N = 3 | n = 12 | p = 0.91 |
| PPX effects on iMSNs (Ctrl) | Fig. 3E | Within unit | WSR | N = 4 | n = 10 | p = 0.38 |
| PPX effects on dMSNs (Park) | Fig. 3F | Within unit | WSR | N = 5 | n = 24 | p = 0.01 |
| PPX effects on iMSNs (Park) | Fig. 3G | Within unit | WSR | N = 4 | n = 16 | p < 0.001 |
| PPX effects on response type  (Ctrl vs. Park) | Fig.  3D&F, insets | Between group | Fisher’s exact | Ctrl: N = 3  Park: N = 5 | Ctrl:  n = 12  Park:  n = 24 | p = 0.01 |
| PPX effects on response type  (Ctrl vs. Park) | Fig.  3E&G, insets | Between group | Fisher’s exact | Ctrl: N = 4  Park: N = 4 | Ctrl:  n = 10  Park:  n = 16 | p = 0.002 |
| PPX effects on response type  (Ctrl vs. Park) | Fig. S4D | Between group | Fisher’s exact | Ctrl: N = 7  Park: N = 11 | Ctrl:  n = 117  Park:  n = 214 | p = 0.0003 |
| Chronic effects of PPX on DD curve (Ctrl) | Fig. 4B | Within animal | WSR, Bonferroni | N = 16 | NA | 0s:  Ctrl vs. 1^st^, p > 0.99  Ctrl vs. 4^th^, p > 0.99  1^st^ vs. 4^th^, p > 0.99  3s:  Ctrl vs. 1^st^, p > 0.99  Ctrl vs. 4^th^, p > 0.99  1^st^ vs. 4^th^, p > 0.99  6s:  Ctrl vs. 1^st^, p > 0.99,  Ctrl vs. 4^th^, p > 0.99  1^st^ vs. 4^th^, p > 0.99  9s:  Ctrl vs. 1^st^, p > 0.99  Ctrl vs. 4^th^, p > 0.99  1^st^ vs. 4^th^, p > 0.99 |
| Chronic effects of PPX on DD curve (Park) | Fig. 4C | Within animal | WSR, Bonferroni | N = 31 | NA | 0s:  Park vs. 1^st^, p < 0.01  Park vs. 4^th^, p < 0.0001  1^st^ vs. 4^th^, p = 0.47  3s:  Park vs. 1^st^, p < 0.05  Park vs. 4^th^, p < 0.0001  1^st^ vs. 4^th^, p < 0.05  6s:  Park vs. 1^st^, p = 0.09,  Park vs. 4^th^, p = 0.08  1^st^ vs. 4^th^, p > 0.99  9s:  Park vs. 1^st^, p = 0.65  Park vs. 4^th^, p < 0.01  1^st^ vs. 4^th^, p < 0.05 |
| PPX effects on response type  1^st^ vs. 4^th^  (Ctrl) | Fig. 4D | Between group | Fisher’s exact | 1^st^: N = 7  4^th^: N = 4 | 1^st^: n = 80  4^th^: n = 27 | p = 0.11 |
| PPX effects on response type  1^st^ vs. 4^th^  (Park) | Fig. 4E | Between group | Fisher’s exact | 1^st^: N = 10  4^th^: N = 8 | 1^st^:n =105  4^th^:n = 75 | p = 0.002 |
| Chronic effects of PPX on ‘increase’  (1^st^ vs. 4^th^) | Fig. 4F | Between group | MWU | 1^st^: N = 6,  4^th^: N = 7 | 1^st^: n = 19  4^th^: n = 32 | p = 0.02 |
| Chronic effects of PPX on ‘decrease’  (1^st^ vs. 4^th^) | Fig. 4G | Between group | MWU | 1^st^:  N = 9  4^th^:  N = 5 | 1^st^:  n = 34  4^th^:  n = 15 | p = 0.89 |

**EXPERIMENTAL MODEL AND SUBJECT DETAILS.**

**Animals**

All mice were on a C57BL/6 background and housed under a 12-h light/dark cycle with ad libitum access to food and water unless stated otherwise. All experiments were performed during the light phase. We used male and female mice aged 3-7 months old. For operant behavior, mice were placed on food restriction 3 days before the onset of magazine training, which was maintained during training and testing phases. Their weight was monitored closely to ensure that their food-restricted body weight remained around 85%-90% of their free-feeding body weight. All experiments were conducted with the approval of the Institutional Animal Care and Use Committee at the University of California, San Francisco, and complied with local and national ethical and legal regulations regarding the use of mice in research.

Hemizygous BAC transgenic mice expressing Cre recombinase under the control of the Drd1a (D1-Cre, GENSAT BAC transgenic EY217) or Adora2a (A2a-Cre, GENSAT BAC transgenic KG139) regulatory elements were used to restrict the expression of Cre-dependent constructs to direct and indirect medium spiny neurons, dMSNs and iMSNs, respectively. For *ex vivo* slice physiology experiments, A2a-Cre mice were bred to hemizygous Drd2-GFP mice, to generate A2a-Cre;Drd2-GFP mice.

**METHOD DETAILS**

**Surgical Procedures**

A detailed surgery protocol can be found at (DOI: [dx.doi.org/10.17504/protocols.io.b9kxr4xn](https://dx.doi.org/10.17504/protocols.io.b9kxr4xn)). Briefly, anesthesia was induced with intraperitoneal (i.p.) injection of ketamine/xylazine (40/10 mg/kg) and maintained with 0.3%-0.8% inhaled isoflurane. Mice were placed in a stereotaxic frame and a mounted drill was used to create holes over the dorsolateral striatum (DLS) or dorsomedial striatum (DMS). To produce parkinsonian or control mice, the bilateral DLS (+ 0.8 mm AP, ± 2.0-2.2 mm ML, − 2.5 mm DV) were injected using a 33-gauge cannula (Plastics One) with 6-hydroxydopamine (6-OHDA)-bromide (1.5 μL per site, 2.5 μg/μL) or saline, respectively. To minimize uptake of the toxin by noradrenergic and serotonergic axons, desipramine (Sigma-Aldrich, 25 mg/kg i.p.) was administered immediately prior to surgery. For subsequent optically identified single-unit recordings, AAV5-DIO-ChR2-eYFP (UPenn Vector Core, 1.0 μL, undiluted) was injected into the left DMS (+ 0.8 mm AP, + 1.5 mm ML, − 2.5 mm DV) in a subset of mice. For chemogenetic inhibition of iMSNs, AAV5-DIO-hM4D(Gi)-mCherry or AAV5-DIO-mCherry (UNC Vector Core, 1.0 μL, diluted 1:2 in normal saline) was injected in bilateral DMS (+ 0.8 mm AP, ± 1.5 mm ML, − 2.5 mm DV). Both 6-OHDA and virus were injected at a rate of 0.2 μL/min, after which the injection cannula was left in place for 10 min prior to being withdrawn, and the scalp being sutured.

In preparation for *in vivo* single-unit recordings, D1-Cre and A2a-Cre mice were injected bilaterally with DLS 6-OHDA (or saline) and DMS DIO-ChR2-eYFP, followed by implantation of DMS optrode arrays in a second surgical procedure. After reopening the scalp, a craniectomy (1.5 x 1 mm) was made over the left DMS. Additionally, two small holes were drilled in the right frontal and right posterior areas for the placement of a skull screw (Fine Scientific Tools, FST) and ground wire, respectively. A fixed multichannel electrode array (32 Tungsten microwires, Innovative Neurophysiology) coupled to a 200 μm optical fiber (Thorlabs) was slowly lowered through the craniectomy into the DLS. The final position of the electrode tips was targeted 150 μm above the previous DIO-ChR-eYFP injection site (-2.35 mm DV). The array was covered and secured into place with dental cement (Metabond) and acrylic (Ortho-Jet). To allow for adequate viral expression, mice were housed for at least two weeks following viral injections before any electrophysiology or behavioral experiments began.

All animals were given buprenorphine (i.p., 0.05 mg/kg) and ketoprofen (subcutaneous injection, 5 mg/kg) for postoperative analgesia. Parkinsonian animals were monitored closely for one week following surgery: their cages were kept on a heating pad, and mice received daily saline injections and were fed nutritional supplements (Diet-Gel Recovery Packs and forage/trail mix).

**Motor assessment**

Details of the accelerating rotarod and open field tests can be found at (DOI: [dx.doi.org/10.17504/protocols.io.q26g7yo4kgwz/v1](https://dx.doi.org/10.17504/protocols.io.q26g7yo4kgwz/v1); DOI: [dx.doi.org/10.17504/protocols.io.b9ksr4we](https://dx.doi.org/10.17504/protocols.io.b9ksr4we)). Briefly, motor function was evaluated using the accelerating rotarod test (Ugo Basile) and open field locomotion at two time points: 3 weeks after 6-OHDA injection surgery and post-pramipexole (PPX) administration. To minimize the effect of motor learning on rotarod performance, each mouse underwent no more than two sessions: one before and one after the PPX administration. Each session included three trials with a 10-min inter-trial interval. The rotation speed gradually increased from 5 to 80 RPM over 5 min. Mice were scored for their latency to fall and the average across three trials was reported. Open field locomotion was also performed in a subset of mice to examine the therapeutic effect of PPX. Mice were habituated to the open field (clear acrylic cylinders, 25 cm in diameter) for 30 min, 1-2 days prior to behavioral sessions. During experimental session, overall movement was monitored with an overhead camera and analyzed offline using video-tracking software (Noldus Ethovision), including distance traveled, velocity, and rotations (90° contralateral or ipsilateral turns).

**Operant Training and Assessment**

A detailed protocol for operant training can be found at (DOI: [dx.doi.org/10.17504/protocols.io.4r3l22k93l1y/v1](https://dx.doi.org/10.17504/protocols.io.4r3l22k93l1y/v1)). Operant behavior was conducted in nine custom-made 18 x 18 x 26 cm operant chambers, enclosed within sound-attenuating cabinets (Coulbourn Instruments). One side of each chamber was equipped with one yellow LED as a house light and remained illuminated during all experimental stages, unless stated otherwise. The opposite side of the chamber was equipped with left and right side nosepokes 12 cm apart. Each nosepoke contained two yellow LED stimulus lights: one positioned 6 cm above and another located inside the nosepoke. Sweetened condensed milk (diluted 1:3 with water) was delivered to a liquid receptacle from the central port, equidistant from the left and right nosepokes through a solenoid valve (The Lee Company). An infrared detector was mounted horizontally across the center port to detect head entries. Experimental events and data collection were managed by a PC running Arduino software. The Arduino scripts for running training and assessment phases are available here: https://zenodo.org/doi/10.5281/zenodo.10703131.

Operant training began around three weeks after 6-OHDA (or saline) injection using a three-phase shaping protocol. Each mouse was trained in the same operant chamber throughout the study. Male and female mice were trained in separate operant chambers. To promote consistency, testing was performed at the same time of day, five days a week.

In Phase 1 (magazine training), which consisted of a single session, food-restricted mice were placed in the operant box, and liquid reward (10 μL of milk) was delivered on a random interval schedule of 40-80 s for a total of 40 rewards. Each trial began with the delivery of milk into the center port, accompanied by a 10 s illumination of the center port light.

In Phase 2 (blocked instrumental learning), mice were trained to nosepoke at the side ports to obtain reward on a fixed-ratio 1 (FR1) schedule. Each training session was separated into two blocks, separated by a two-minute break period when the house light was turned off. Each block lasted 50 min. Only one nosepoke (either the left or the right side) was trained in each block, and the active nosepoke switched during the break period. The initial nosepoke (left or right) was counterbalanced across sessions. Each trial began with a stimulus cue light above the left or right nosepoke being illuminated. If the mouse nosepoked that port within 20 s, the cue light was extinguished, and reward (10 μL of milk) was delivered to the center port, which was illuminated for 10 s, followed by a random 30-50 s inter-trial interval. If the mouse did not perform a nosepoke on the correct side, then cue lights were extinguished and went into a timeout (a random 30-50 s interval). Mice were trained in Phase 2 for at least 6 sessions and until they met criteria (correct response on > 80% of trials). Mice that did not meet criteria after 10 sessions were removed from the study. Overall, 6 out of 82 mice were excluded during Phase 2.

Phase 3 (randomized instrumental learning) resembled Phase 2 with two additions: the introduction of self-initiated trial start, and pseudorandomized (no more than two consecutive trials with the same side cued), rather than blocked trials. Trials began with the center port being illuminated for 10 s. During this period, a trial could be initiated by a center port nosepoke. Failure to make a center nosepoke within this period was considered a center omission and was followed by a timeout period (random 30-50 s interval). Following an initiation nosepoke, the stimulus cue lights at either the left or right port were activated for 10 s. If the mouse nosepoked that port within 10 s, the cue light was extinguished, and reward (10 μL of milk) was delivered to the center port, which was illuminated for 10 s, followed by a random 30-50 s inter-trial interval. Failure to poke the correct side port within 10 s was considered a side omission, and cue lights were extinguished until the next trial. Poking the opposite, uncued side was recorded as an incorrect response, but did not trigger punishment. Each session was 90 min. Mice were trained in Phase 3 for at least 5 sessions and until criteria were met. Criteria were (1) responding correctly in at least 80% of trials, and (2) incorrect responses were less than 15% of the total (to reduce side bias). Mice that did not meet criteria in 8 sessions were excluded. Overall, 5 out of 76 mice were excluded during Phase 3.

Phase 4 (delay discounting) was the final task. The delay discounting task was composed of three trial types: forced choice delayed/large, forced choice immediate/small, and free choice. Each session was composed of four blocks. Each block included 30 trials, starting with five forced choice delayed/large trials, followed by five forced choice immediate/small trials to remind mice of the delay contingencies in effect for that block. These “refresher” trials were followed by 20 free choice trials. In a given mouse, the left and right sides were assigned to either delayed/large or immediate/small outcomes, but the side assignments were randomly distributed between mice. Each trial was a fixed duration of 50 s. Therefore, port choice did not influence the trial duration (i.e., choosing the small/immediate reward did not lead to the next trial quicker). All trials started with 10 s illumination of the center port light. A center nosepoke within 10 s extinguished the light and initiated the trial. Trials on which mice failed to nosepoke during this window were recorded as center omissions. During forced choice delayed/large trials, cue lights on one side were illuminated for 10 s, and nosepoke at that port resulted in a large reward (15 μL) delivered to the center port after a delay period, as described below. During forced choice immediate/small trials, cue lights on the other side were illuminated for 10 s, and nosepoke at that port resulted in a small (5 μL) reward delivered to the center port immediately. During free choice trials, both left and right cue lights were illuminated for 10 s. Nosepokes on either side resulted in reward, following the contingencies introduced during the “refresher” trials. The delay preceding large reward delivery increased across a session (between blocks) from 0 s (block 1) to 3 s (block 2), to 6 s (block 3) to 9 s (block 4). If a mouse failed to nosepoke in one of the side ports within 10 s, cue lights were extinguished, and the trial was recorded as a side omission. After reward delivery or side omission, the inter-trial interval (adjusted to achieve trial start every 50s) began. After 2-3 weeks of training on the task, once mice had established a stable delay discounting curve (see the criteria in the experimental design section), the delay discounting performance was evaluated in “pramipexole (PPX) ON” (4h after PPX), and “OFF” (48h after PPX) states. Overall, 11 out of 69 mice were exclude in Phase 4 from the study because their delay discounting curves were not established and stable within 3 weeks.

***In vivo* Electrophysiology**

A detailed protocol for *in vivo* electrophysiology can be found at (DOI: [dx.doi.org/10.17504/protocols.io.b9ucr6sw](https://dx.doi.org/10.17504/protocols.io.b9ucr6sw)). Briefly, one week after the optrode array implantation, mice were habituated to tethering and the open field chamber for at least 2 days. After habituation, experimental sessions occurred at least once per week for 4-6 weeks. During each session, animals were plugged into a lightweight multiplexed and commutated headstatge cable to record single-unit activity (CerePlex Direct, Blackrock Microsystems). Spike waveforms were filtered at 154-8800 Hz and digitized at 30 kHz. The experimenter manually set a threshold for storage of electrical events. Spike sorting and single units were identified offline by manual sorting into clusters (Offline Sorter, Plexon). Waveform features used for separating units were typically a combination of valley amplitude, the first three principal components (PCs), and/or nonlinear energy. Clusters were classified as single units if they fulfilled the following criteria: (1) the unit’s waveforms were statistically different from multiunit activity and any other single units on the same wire, in 3D PCA space, (2) no inter-spike interval < 1 msec was observed. Single-units were then classified as putative medium spiny neurons (MSNs) or interneurons as previously described ^29,30^ using features of the spike waveform (peak to valley and peak width), as well as inter-spike interval distribution. After single-units had been selected for further study, their firing activity was analyzed using NeuroExplorer 4.133 (Nex Technologies). To determine if a unit was optogenetically identified, a peristimulus time histogram was constructed around the onset of laser pulses. To be considered optogenetically identified, a unit had to fulfill 3 criteria: (1) the unit had to increase firing rate above the 99% confidence interval of the baseline within 15 msec of laser onset; (2) the unit’s firing was above this threshold for at least 15 msec; (3) the unit’s laser-activated waveforms were not statistically distinguishable from spontaneous waveforms.

During *in vivo* single-unit recordings, mice were injected i.p. with either PPX or saline, after which their gross locomotion was monitored with video-tracking of locomotor activity for 330 min (see details in the experimental design section).

***Ex vivo* Electrophysiology**

A detailed protocol can be found at (DOI: [dx.doi.org/10.17504/protocols.io.b9uir6ue](https://dx.doi.org/10.17504/protocols.io.b9uir6ue)). Briefly, slice electrophysiology was used to validate the inhibitory Designer Receptor Exclusively Activated by Designer Drug (DREADD) for chemogenetic inhibition experiments. Acute slices from A2a-Cre;Drd2-GFP mice coinjected with AAV5-DIO-ChR2-eYFP (UPenn Vector Core) and AAV5-DIO-hM4D(Gi)-mCherry (UNC Vector Core) were prepared. Mice were deeply anesthetized with ketamine-xylazine (100-200 mg, i.p.) and perfused with a carbogenated, ice-cold glycerol-based artificial cereberospinal fluid (ACSF) solution containing (in mM): 250 glycerol, 2.5 KCl, 1.2 NaH_2_PO_4_, 10 HEPES, 21 NaHCO_3_, 5 D-glucose, 2 MgCl_2_, 2 CaCl_2_. Following decapitation, brains were dissected, mounted on a chuck, and submerged in ice-cold glycerol solution. A vibrating microtome (Leica) was used to cut sequential 275 mm coronal slices containing the striatum, which were immediately transferred to warm (34°C), carbogenated ACSF containing (in mM): 125 NaCl, 26 NaHCO_3_, 2.5 KCl, 1.25 NaH_2_PO_4_, 12.5 D-glucose, 1 MgCl_2_, 2 CaCl_2_. Slices were incubated for 30-60 min, then kept at room temperature until use.

During all recordings, slices were superfused with carbogenated ACSF at 31-33°C. Differential interference contrast (DIC) optics on an Olympus BX 51 WIF microscope were used to target MSNs, which were patched in a whole-cell configuration using borosilicate glass electrodes (2-5 MΩ). Whole-cell voltage-clamp recordings were made using a MultiClamp 700B amplifier (Molecular Devices) and ITC-18 A/D board (HEKA). Data was acquired using Igor Pro 6.0 software (Wavemetrics) and custom acquisition routines (mafPC, courtesy of M. A. Xu-Friedman). Recordings were filtered at 2 kHz and digitized at 10 kHz. To measure the acute effect of CNO on the synaptic output of iMSNs in slice, dMSNs were targeted for whole-cell recordings and identified by their GFP/eYFP-negative somata in striatal regions showing mCherry positive processes. GFP/eYFP-negative neurons with GABAergic interneuron physiological properties (membrane tau decay < 1 ms) were excluded from further analysis. All synaptic currents were recorded with a cesium methanesulfonate-based internal with high chloride, which contained (in mM): 120 CsCl, 15 CsMESO_3_, 8 NaCl, 0.5 EGTA, 10 HEPES, pH = 7.3 and monitored at a holding potential of -70 mV. Series resistance and leak currents were monitored continuously. Inhibitory synaptic currents were optically evoked using 3 ms pulses of 473 nm light ranging in power from 0.5-4 mW and delivered by a TTL-controlled LED (Olympus) passed through a GFP filter (Chroma). oIPSCs were elicited every 20 s and the amplitudes were compared before and 10-15min after addition of CNO to the ACSF (1 μM). This slice validation data for chemogenetic inhibition was also utilized in another published work^31^.

**Pharmacology**

6-OHDA (Sigma Aldrich) for striatal dopamine depletions was prepared at 2.5 μg/μL in normal saline on the day of surgery and wrapped in aluminum foil until use. Pramipexole dihydrochloride (Sigma Aldrich) was prepared in normal saline solution and administrated via i.p. injection at a dose of 0.5 mg/kg, at least four times during the delay discounting assessment stage. Cognitive or motor assessments were conducted approximately 4 h after PPX injection. At this time point, locomotion was increased (avoid the initial period of reduced locomotion) ^32^. Clozapine-N-oxide (CNO, Tocris Bioscience) was dissolved in normal saline at 0.3 mg/ml, and injected i.p. at a final dose of 3 mg/kg. This CNO concentration was chosen as it increased locomotion in parkinsonian mice. The stock solution of CNO was wrapped in aluminum foil to minimize light exposure. For *ex vivo* experiments, CNO was dissolved in normal saline and then diluted in ACSF for a final concentration of 1 μM.

**Histology, Quantification & Microscopy**

A detailed protocol for preparation of histological sections can be found at (DOI: [dx.doi.org/10.17504/protocols.io.b9ubr6sn](https://dx.doi.org/10.17504/protocols.io.b9ubr6sn)). After behavioral, *in vivo* electrophysiology, and chemogenetic experiments, mice were deeply anesthetized with IP ketamine-xylazine and transcardially perfused with 4% paraformaldehyde in PBS. Following *in vivo* electrophysiology experiments, prior to perfusion, electrode array location was marked by electrolytic lesioning. After deep anesthesia, the implant was connected to a solid state, direct current (DC) Lesion Maker (Ugo Basile). A current of 100 μA was passed through each microwire for 5 s. After perfusion, the brain was dissected from the skull and post-fixed overnight in 4% paraformaldehyde, then placed in 30% sucrose at 4°C for cryoprotection. The brain was then cut into 30 μm coronal sections on a freezing microtome (Leica) and then mounted in Vectashield Mounting Medium onto glass slides for imaging. For immunohistochemistry, the tissue was blocked with 3% normal donkey serum (NDS) and permeabilized with 0.1% Triton X-100 for 2 h at room temperature on a shaker. Primary antibodies were added to 3% NDS and incubated for 2 days at 4°C on a shaker. Primary antibodies used: Rabbit anti-TH (Pel-Freez, 1:1000), Chicken anti-TH (Sigma, 1:1000). Slices were then incubated in secondary antibodies (donkey anti-rabbit or chicken Alexa fluor 488 or 647, 1:500, JacksonImmuno Research) overnight at 4°C on a shaker, washed, and mounted onto slides for imaging. Images at 10x magnification for striatal sections and 20x for SNc sections were acquired using a Nikon 6D conventional widefield microscope. Consistent lighting and exposure settings were maintained for all sections from both healthy and parkinsonian groups to allow for reliable comparison and subsequent image analysis. The Mouse Brain Atlas in Stereotaxic Coordinates (hard copy, 4^th^ edition) was consulted for anatomical reference.

To quantify the residual TH positive fluorescence intensity within the striatum, 5-8 coronal sections between (AP 0.38 mm-1.18 mm) were analyzed. Regions of interest (ROIs) were outlined using the freehand ROI tool in ImageJ based on anatomical landmarks for striatum. The ROIs included 3 subregions: dorsolateral striatum (DLS), dorsomedial striatum (DMS), and ventral striatum (VS). Subsequent analysis involved setting measurement parameters to include Area, Raw Integrated Density, and Mean Gray Value. These parameters were measured for each ROI and 4 background areas, which showed minimal fluorescence to account for background signal. The mean background fluorescence was first calculated, followed by the calculation of corrected total fluorescence (CTF) using the formula: (Raw Integrated Density-(Area x Mean Fluorescence of background)) / Area of ROI.

To quantify the residual dopaminergic neurons within the midbrain, the substantia nigra pars compacta (SNc) region was outlined using the Allen Brain Atlas as a reference. TH-positive cells were manually counted per section using the point tool function in ImageJ. Counting was performed bilaterally and blinded to experimental condition across 4 stained sections per mouse, (AP -3.07 mm, -3.15 mm, -3.39 mm and -3.51 mm). Dopaminergic cell density was calculated and compared between healthy and parkinsonian conditions.

**EXPERIMENTAL DESIGN AND STATISTICAL ANALYSIS**

The experimental design and statistical analysis of all key experiments are summarized in Table 1. This table includes statistical tests, N (animals), n (units or cells), p values and the associated figures. All data are presented as the mean ± SEM. The statistical tests were performed using GraphPad Prism 10. Since results sometimes violate the assumption of normality of residuals as assessed by Shapiro-Wilk test, the effects of PPX on delayed/large reward choice, omissions, trigger latency were compared within subjects using multiple Wilcoxon Signed-Rank test, and Wilcoxon Rank-Sum tests (denoted Mann-Whitney U test) between groups. In all analyses, a p value of < 0.05 was considered statistically significant, unless Bonferroni-corrected for multiple comparisons (detailed below), in which case the p value was multiplied by the number of comparisons.

*Behavior*

For analysis of delay discounting performance, the primary outcome was the percentage of large reward choices at each delay. Secondary outcome measures included other parameters (detailed below) that characterize overall delay discounting behavior during free choice trials. Raw data files were analyzed using custom analysis code written in Python (https://zenodo.org/doi/10.5281/zenodo.10703139). This code extracted specific task-related events, including trial start, cue light start, left and right nosepokes, reward consumptions and timeouts during both forced and free choice trials. Choice performance in each block of the delay discounting task was measured as the percentage of free choice trials (excluding omissions) on which mice chose the delayed/large reward. Each mouse was trained on the delay discounting task until it reached stable baseline performance. Stable baseline was first assessed in each mouse by visually inspecting discounting curves for three sessions, then by calculation of the coefficient of variation for choice of the delayed/large outcome (stable defined as CV < 20% in at least 3 blocks for three consecutive sessions). Once this criterion was met, delay discounting behavior was tested in the following conditions: baseline (pretreatment); PPX ON (4 h following the 1^st^, 2^nd^, 3^rd^, and 4^th^ PPX injections); and PPX OFF (48 h after each PPX injection, with a saline injection given instead). Latencies to choose delayed/large rewards and immediate/small rewards during free choice trials were measured as the interval between the illumination of the nosepoke (side) cue light and a nosepoke (side) response, excluding side omissions. The effect of PPX on the delay discounting performance, including delayed/large reward choice, latency to choose, and omissions, at different delays, were determined using the multiple Wilcoxon Signed-Rank test with Bonferroni correction between baseline and PPX ON or PPX OFF conditions. To measure the chronic effects of PPX on the choice of delayed/large reward during delay discounting task, the 1^st^ and 4^th^ PPX injections were compared across different delays using multiple Wilcoxon Signed-Rank test with Bonferroni correction.

To better understand the effects of PPX on delay discounting performance in which PPX significantly favored small/immediate choice in parkinsonian mice, additional analyses were conducted to capture how PPX specifically altered the shape of the discounting curve. The data was fitted to Herrnstein’s simple hyperbolic model: V = *A*/(1+*K*D) to estimate the discount rate parameter (*K)*, reward size parameter (*A*), and the area under the fitted curve (AUC) ^33,34^. Effects of PPX on these parameters in parkinsonian group were analyzed using Wilcoxon Signed-Rank test.

For assessing motor ability, the primary outcome measures were the latency to fall (accelerating rotarod test) and average velocity (open field test), compared between healthy and parkinsonian mice using the Mann-Whitney U test test. The therapeutic effects of PPX on rotarod performance and velocity (240-300 min after PPX) were determined using the Wilcoxon Signed-Rank test.

*In Vivo Electrophysiology*

*In vivo* electrophysiology sessions lasted 330 min. Firing rates were averaged in 1 minute bins. The modulation of firing rate by PPX was determined by comparing average firing rate before (-30-0 min) and after (240-300 min) PPX administration. This epoch matched the timing of the cognitive assessments after PPX. The 30-min baseline period was compared to two 30-min periods following drug injection (241-270 min and 271-300 min post-injection). Both optically labeled and unidentified MSNs were categorized into three groups as follows, based on the direction of change in firing rate according to Wilcoxon Signed-Rank test (*p* < 0.01): increase type (significant increase in both 30-min post injection periods), decrease type (significant decrease in both 30-min post injection periods) or no change units (no significant change in either of 30-min post injection periods).

Fisher's exact test was performed to compare the percentages of the three response types (increase, decrease and no change) between the 1^st^ and 4^th^ PPX injections in both control and parkinsonian groups. Firing rates of optogenetically labeled dMSNs and iMSNs recorded from parkinsonian mice were compared to recordings from healthy mice using the Mann-Whitney U test. Firing rates of optically-identified MSNs in parkinsonian mice before and after PPX administration were compared using the Wilcoxon Signed-Rank test.

*Chemogenetic inhibition*

In inhibitory DREADD validation slice electrophysiological experiments, oIPSC amplitudes were compared before and 10-15 min after addition of CNO, using the Wilcoxon Signed-Rank test. The effect of chemogenetic inhibition on gross locomotion was assessed by comparing average movement velocity during baseline (-20-0 min before CNO) and 20-30 min following CNO administration, using the Wilcoxon Signed-Rank test. In chemogenetic experiments during the delay discounting task, performance was compared within subjects in pre-CNO treatment periods versus 30 min post-CNO administration. Similar to PPX administration experiments, a stable baseline was established across at least three consecutive sessions. Post CNO administration sessions were averaged per animal across four CNO treatment sessions. Performance in free choice delay discounting trials across different delays were compared using the multiple Wilcoxon Signed-Rank test with Bonferroni correction. Other delay discounting parameters (*A*, *K* and AUC) were also compared in pre- and post-CNO sessions using the Wilcoxon Signed-Ranks test.

*Quantitative Histology*

The relationship between dopamine depletion and delay discounting behavior parameters (*A*, *K* and AUC*)* after PPX administration was quantified with Pearson correlation. Correlation coefficient (r) and p values were used to describe the linear correlation, and the coefficient of determination r^2^ was used to describe the power of the model.

| KEY RESOURCES TABLE | | |
| --- | --- | --- |
| Reagent or Resource | Source | Identifier |
| Antibodies | | |
| Rabbit anti-TH | Pel-Freez Biologicals | Cat# P40101-150, RRID: AB_2617184 |
| Chicken anti-TH | Millipore | Cat# AB9702, RRID: AB_570923 |
| Alexa Fluor 488-Donkey Anti-Rabbit IgG | Jackson ImmunoResearch Labs | Cat# 711-546-152, RRID: AB_2340619 |
| Alexa Fluor 647-Donkey Anti-Rabbit IgG | Jackson ImmunoResearch Labs | Cat# 711-606-152, RRID: AB_2340625 |
| Alexa Fluor 647-Donkey Anti-Chicken IgY (IgG) | Jackson ImmunoResearch Labs | Cat# 703-606-155, RRID: AB_2340380 |
| Virus Strains | | |
| AAV5-EF1a-DIO-hCHR2(H134R)-eYFP-wpre-hGH | Penn Vector Core | Lot #CS1046  Addgene viral prep # 20298-AAV5; http://n2t.net/addgene:20298; RRID:Addgene_20298 |
| AAV5-hSyn-DIO-hM4D(Gi)-mCherry | UNC | Lot #V120251  (Addgene viral prep # 44362-AAV5; http://n2t.net/addgene:44362; RRID:Addgene_44362 |
| AAV5-hSyn-DIO-mCherry | UNC | Lot #AV4634C  Addgene viral prep # 50459-AAV5; http://n2t.net/addgene:50459 ; RRID:Addgene_50459) |
| Chemicals | | |
| 6-Hydroxydopamine hydrobromide | Sigma-Aldrich | 162957 |
| Pramipexole dihydrochloride | Sigma-Aldrich | A1237 |
| Desipramine hydrochloride | Sigma-Aldrich | D3900 |
| Clozapine-N-oxide | Tocris Bioscience | Cat# 6329 |
| Critical Commercial Assays | | |
| VECTASHIELD Antifade Mounting Medium | Vector Laboratories | Cat# H-1000, RRID: AB_2336789 |
| Experimental Models: Organisms/Strains | | |
| Mouse: WT: C57BL/6J | The Jackson Laboratory | RRID: IMSR_JAX:000664 |
| Mouse: B6.FVB(Cg)-Tg (Drd1-cre) EY217Gsat/ Mmucd | MMRRC | RRID: MMRRC_034258-UCD |
| Mouse: B6.FVB(Cg)-Tg (Adora2a-cre) KG139Gsat/ Mmucd | MMRRC | RRID: MMRRC:036158-UCD |
| Mouse: STOCK Tg(Drd2-EGFP)S118Gsat/ Mmnc Mus musculus | MMRRC | RRID: MMRRC_000230-UNC |
| Software and Algorithms | | |
| Igor Pro | Wavemetrics | http://www.wavemetrics.com/products/igorpro/ igorpro.htm; RRID: SCR_000325  With MafPC:  https://www.xufriedman.org/mafpc |
| MATLAB R2020b | MathWorks | https://www.mathworks.com/products/matlab.html; RRID: SCR_001622 |
| EthoVision XT | Noldus | https://www.noldus.com/ethovision-xt;  RRID: SCR_000441 |
| Adobe Illustrator CS5 | Adobe | https://www.adobe.com/products/illustrator.html;  RRID: SCR_014198 |
| MAP Software | Plexon | https://plexon.com/products/map-software;  RRID: SCR_003170 |
| Offline Sorter | Plexon | https://plexon.com/products/offline-sorter;  RRID: SCR_000012 |
| NeuroExplorer | Nex Technologies | http://www.neuroexplorer.com/;  RRID: SCR_001818 |
| GraphPad Prism | GraphPad | https://www.graphpad.com/features;  RRID: SCR_002798 |
| Other | | |
| 32-channel fixed optrode array | Innovative Neurophysiology | Custom; http://www.inphysiology.com/ optogenetic-applications/ |
| 200 μm Core TECS-Clad Multimode Optical Fiber, 0.39 NA | Thorlabs | Cat# FT200UMT |
| 1.25 mm Multimode LC/PC Ceramic Ferrule,230 mm Bore Size | Thorlabs | Cat# CFLC230-10 |
| 150mW DPSS 473nm Blue Laser | Shanghai Laser & Optics Century | BL473T8-150 + ADR-800A |
| 200 mm Core, 0.39 NA FC/PC to Ø1.25 mm Ferrule Patch Cable, 1 m Long | Thorlabs | Cat# M83L01 |
| Blackrock Acquisition System | CerePlex Direct,  Blackrock Microsystems | https://blackrockneurotech.com/support/ |
